# Multi-site assessment of reproducibility in high-content live cell imaging data

**DOI:** 10.1101/2022.11.18.516878

**Authors:** Jianjiang Hu, Xavier Serra-Picamal, Gert-Jan Bakker, Marleen Van Troys, Sabina Winograd-katz, Nil Ege, Xiaowei Gong, Yuliia Didan, Inna Grosheva, Omer Polansky, Karima Bakkali, Evelien Van Hamme, Merijn Van Erp, Manon Vullings, Felix Weiss, Jarama Clucas, Anna M. Dowbaj, Erik Sahai, Christophe Ampe, Benjamin Geiger, Peter Friedl, Matteo Bottai, Staffan Strömblad

**Affiliations:** Department of Biosciences and Nutrition, Karolinska Institutet, Stockholm, Sweden; Department of Cell Biology, Radboud University Medical Center, Nijmegen, The Netherlands; Department of Biomolecular Medicine, Ghent University, Ghent, Belgium; Department of Immunology and Regenerative Biology, Weizmann Institute of Science, Rehovot, Israel; The Francis Crick Institute, London, United Kingdom; Bio Imaging Core, VIB Center for Inflammation Research, Ghent, Belgium; Division of Biostatistics, Institute of Environmental Medicine, Karolinska Institutet, Stockholm, Sweden

**Author notes:** Equal contribution (Jianjiang Hu, Xavier Serra-Picamal).

## Abstract

High-content image-based cell phenotyping provides fundamental insights in a broad variety of life science areas. Striving for accurate conclusions and meaningful impact demands high reproducibility standards, even more importantly with the advent of data sharing initiatives. However, the sources and degree of biological and technical variability, and thus the reproducibility and usefulness of meta-analysis of results from live-cell microscopy have not been systematically investigated. Here, using high content data describing features of cell migration and morphology, we determine the sources of variability across different scales, including between laboratories, persons, experiments, technical repeats, cells and time points. Significant technical variability occurred between laboratories, providing low value to direct meta-analysis on the data from different laboratories. However, batch effect removal markedly improved the possibility to combine image-based datasets of perturbation experiments. Thus, reproducible quantitative high-content cell image data and meta-analysis depend on standardized procedures and batch correction applied to studies of perturbation effects.

## Introduction

High content cell imaging enables great advances in many life sciences fields, such as cell biology, biomedicine and drug development. Modern microscope setups can generate vast amounts of high resolution data, rich across multiple dimensions, including high spatial and temporal resolution, to differentiate cell structures in a multiplex manner and to spatially resolve and quantify gene or protein expression, as well as the effects of drug perturbation^1, 2^. Accompanying these technological advances, initiatives have emerged to host and make image-based datasets publicly available to the research community, including but not limited to the Image Data Resource, the Cell Image Library and the Human Cell Atlas^3-6^. These platforms have improved the standards for data reporting, with more transparent datasets made available in a sustainable manner^7^. However, to further consolidate reproducible microscopy research, retrieving and cross-correlating image data accessible from different laboratories is required to reuse the data for secondary purposes and to perform meta-analysis studies. An obstacle to this is that we so far lack guidelines and rules for implementation and reuse of high-content imaging data from different sources and, arguably, variability in procedures. Consequently, the data variability between laboratories typically lack standardization and are not suitable for high-quality meta-analysis studies^8^.

Other types of complex data in the life sciences have for long been shared and extensively reused. As examples, multiple studies have addressed the reproducibility of data produced by different laboratories, for instance of mass spectrometry and RNA-seq based data^9-12^.

With the aim of building an open data ecosystem for cell biology research through standardization, dissemination and meta-analysis efforts, the Multimot consortium was established to develop concrete standards for high-quality cell migration research^13-16^. Here, we present a study by five laboratories of the Multimot consortium, where we quantified the sources of variability at different scales in high content imaging data of migrating cancer cells in 2D and 3D environments. Importantly, the highest technical variability occurred between laboratories, preventing direct high-quality meta-analysis of the primary data. However, in perturbation experiments, the variability could be overcome by a batch effect removal approach to achieve reliable meta-analyses of imaging-based datasets from different sources.

## Results

### 2D live cell imaging design and performance

To quantify the sources of variability, a live cell imaging design of cell migration on a 2D surface was replicated in a multi-level, nested structure. Migration behavior of HT1080 fibrosarcoma cells, stably expressing LifeAct-mCherry and H2B-EGFP, on a collagen coated glass surface was recorded using automated fluorescent light microscopes equipped with an environmental chamber. A detailed common protocol (Supplementary material 2-4) was designed and distributed to all participating laboratories as well as the cell line and all key reagents, aiming at minimizing the biological and technical variance. The design involved three independent laboratories, three persons at each laboratory, three independent experiments by each person, two conditions (control and ROCK inhibitor) in each experiment, and three technical replicates in each condition (Fig. 1a). In each technical replicate, around 50 cells were imaged in 5 min intervals for 6 h (Fig. 1b). Experiments were carried out independently by the three participating laboratories, and deviations from the original protocol were kept for the record, including independent microscopy platforms, objective specifications, control hardware for climatization of the cell cultures during microscopy, reagent differences, as well as how strictly the protocol was followed (Supplementary material, the Excel tables). All microscope-derived image collection, data processing and statistical analysis was conducted by the Strömblad laboratory. The uniform data analysis secures identical post experiment data processing and allows to uncover sources of variability in the experimental procedures.

**Figure 1.**
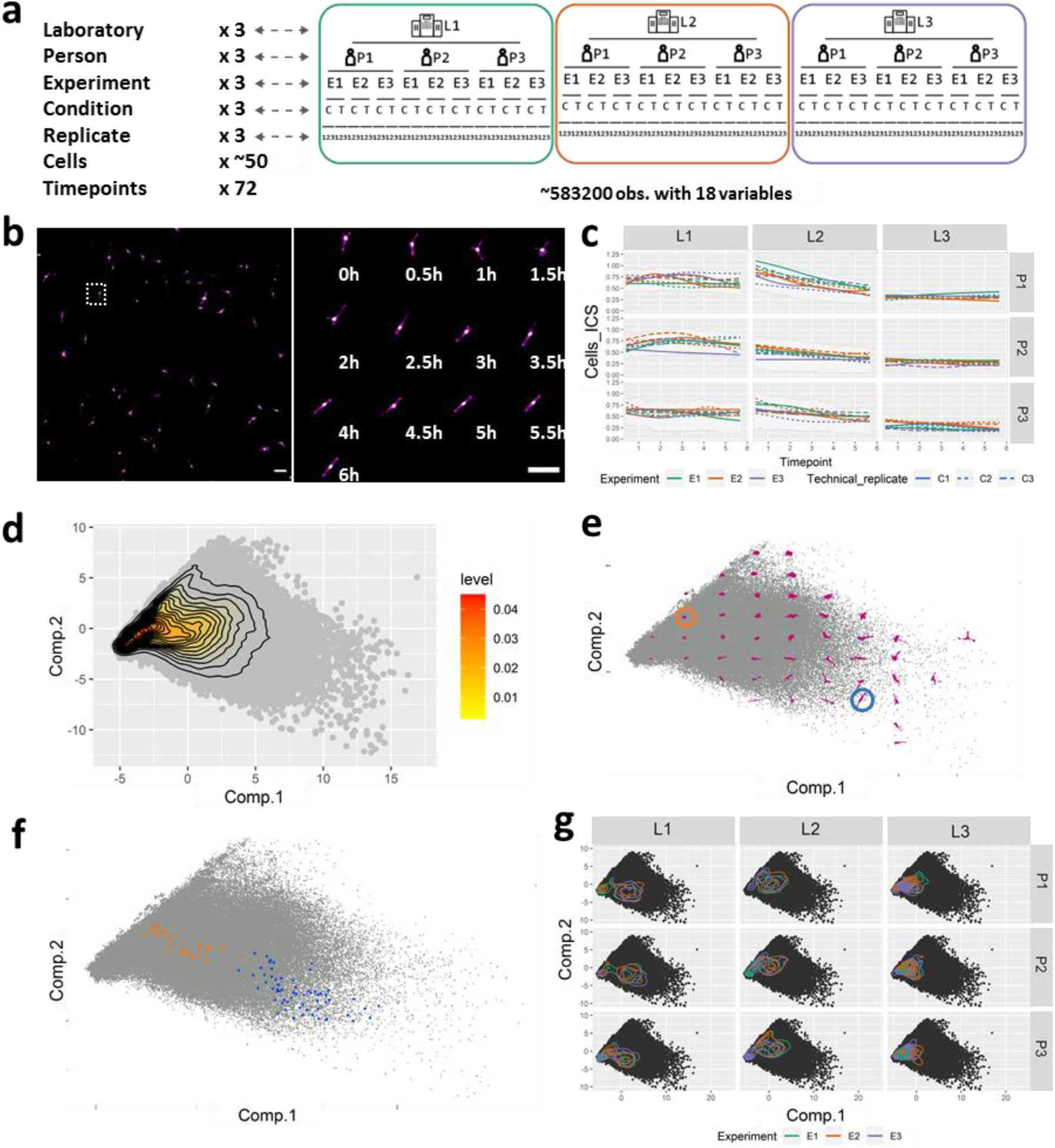
Study design and initial results. **a. Schematic of the study design**. The study involved three independent laboratories, three persons in each laboratory, three independent experiments by each person, two conditions (control or ROCK inhibitor) in each experiment, and three replicates in each condition. For each replicate, around 50 cells were imaged for 6 h in 5 min time intervals. 18 variables were quantified from each image series. **b. Example of acquired time lapse images. Left**: stitched large image; **right**: cropped images of one cell at different time points. Scale bar: 100 μm **c. Quantification results of Instantaneous Cell Speed (ICS) over time for each lab (L1-3), person (P1-3), experiment (E1-3), and technical replicate (C1-C3)**. Lines in different colour represent the data from the control condition (untreated cells) of three different experiments. Different style of the lines with the same colour represent three different technical replicates within one experiment. The error bar indicates the first and third quartiles of the data at each time point. **d. Principal component analysis results of all variables extracted from the entire data**. Grey dots show the position of the first and second principal components for each observation. Inset marks the density of the observation dots. **e. Visualization of cell shapes at different locations of the PCA space**. Grey dots show the position of the first and second principal components for each observation. Representative cell shapes at specific locations in the PCA plot are shown in magenta. **f. The locations of the same cell at different time points within the PCA plot**. Grey dots show the position of the first and second principal components for each observation. Orange and blue dots show the locations of two different cells (dash circled in **e**) in the PCA space at different time points. **g. Principal component analysis results shown for each person (P) in each laboratory (L)**. Black dots show the position of the first and second principal components for each observation from the control condition (untreated cells). Colored lines show the 2D density plots of the technical replicates, where lines with different colors in the same plot represent different experiments. The principal component space is identical in all the plots.

### Data description

For all image time sequences, cellular and nuclear variables were automatically extracted using CellProfiler by the same cell segmentation and tracking strategy, followed by Matlab processing to define protrusion, retraction, and short lived cell regions^17^ based on the CellProfiler derived cell masks (see Methods section for details). The raw images, CellProfiler pipeline, and Matlab scripts have been shared in the SciLifeLab Data Depository. As a result, a total of 18 variables describing either morphological or dynamic features of the cell or the nucleus were obtained and further analyzed. Results accounted for the evolution of each variable over time, for each laboratory, person, experiment, technical replicate and cell (Fig 1c, Supplementary figure 1), were displayed to identify differences in the magnitude or trends of the described variables at these different levels.

Z-score standardization was applied to all features, and subsequent principal component analysis (PCA) was performed in order to maximize and visualize the variability. The first two principal components represent >60% of the variability in the observations (Supplementary figure 2). By combining all observations, we found that the data concentrate around the mean value and dissipate progressively from there, without apparent differentiated clustering of observations in the PCA space (Fig. 1d) ^18^. Observations with different cell shape or the same cell at different time points locate at different places of the PCA space (Fig. 1e-f). Differences in data localization, variability, and clustering were detectable by 2D principal component analysis representing variations among technical repeats, experiments, persons or labs (Fig 1g, Supplementary figure 3).

### Variability sources

We next quantified this variability across the different levels of the hierarchical experiment structure. For this, we modelled the data using Linear Mixed Effect (LME) model for random effects. To identify the sources of variabilities at different levels, we applied the LME model to the control experiments for each of the 18 obtained variables, as well as for the first and second principal components. From the model, we obtained the variance components at each of the levels (temporal, cell, technical replicate, experiment, person, and laboratory) for each variable (Supplementary figure 4a, b)) and categorized the sources of variability as biological or technical variability. Biological variability originated from the cell identity (cells in a population display variability for a given variable) and temporal variation (the same cell displays variability for a given variable when studied at different time points). Technical variability originated from the technical replicate, experiment, person, and laboratory. There was substantial biological variability within the cell population and for each cell over time (Supplementary figure 4a-c). By aggregating the variabilities, we identified technical sources to contribute 32% (median value) of the total variance across all variables (Supplementary figure 4d). While proper study design in terms of the sample size (number of cells, etc.) should take the inherent biological variability into account to facilitate the detection of statistically discernable differences, the reproducibility of the data is defined by their technical variability. Importantly, among the technical variability, lab to lab variability was the major source of variability, followed by person, experiment and replicate, but with different relative contributions among different variables (Fig. 2a-b, Supplementary figure 4c).

**Figure 2.**
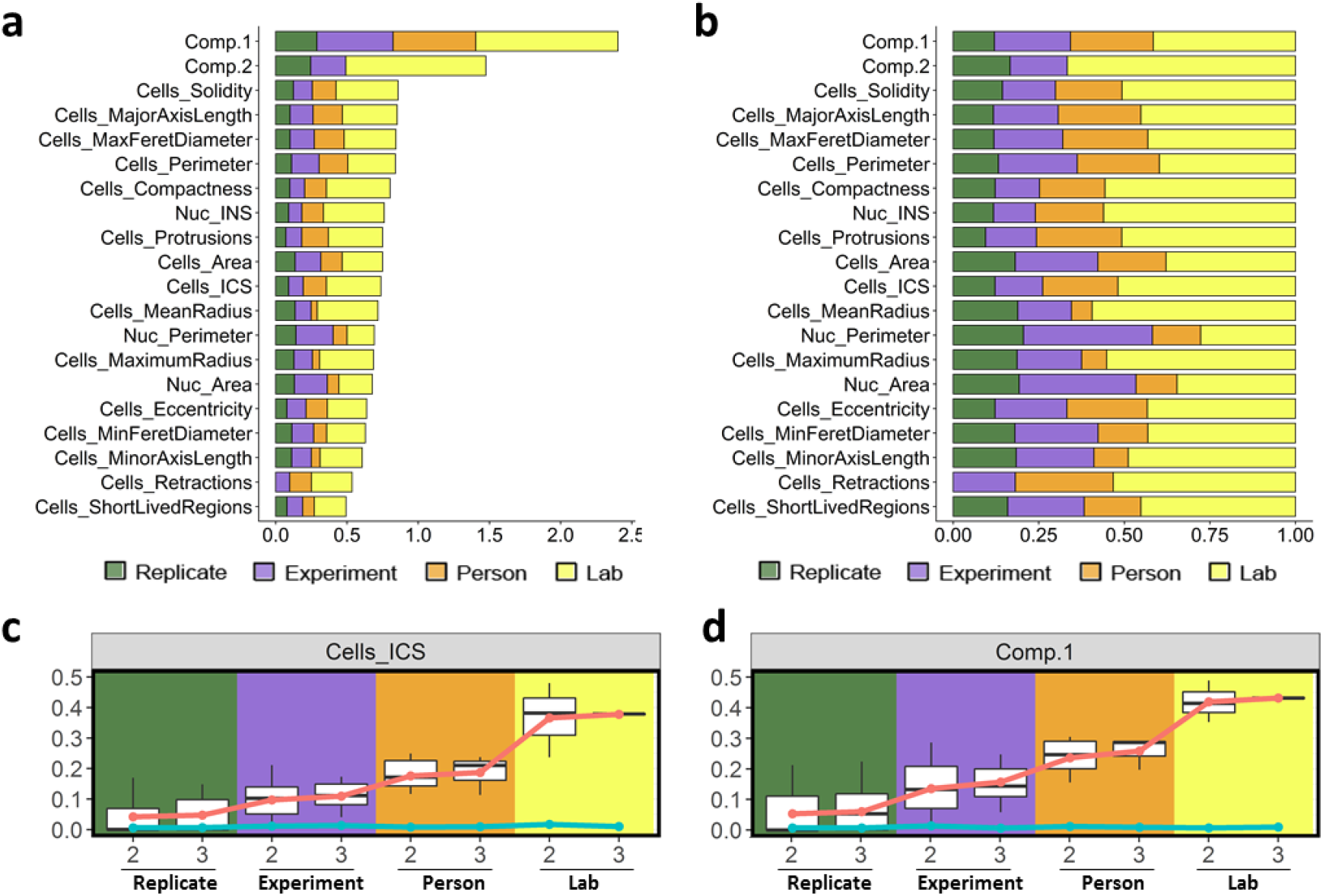
Lab to lab variance contributes the most to the technical variance. **a-b. Variance components of each variable from all technical levels based on the Linear Mixed Effect (LME) model analysis. a:** absolute value; **b:** relative value. **c-d. Cumulative variability of Instantaneous Cell Speed (ICS) (c) and first principal component (d) at the levels of technical replicate, experiment, person, and laboratory**. Boxplots show variances with 2 or 3 replicates, experiments, persons, or laboratories, calculated at each level. Red dots show the mean value of the cumulative variance that are linked with red lines. As a control, cyan dots and lines show the cumulative variance of the same data after randomization.

We then determined the source of technical variability in more detail at each level. We computed the cumulative variability deriving from technical sources when adding additional levels to a hypothetical experimental design with increasing complexity (Methods and Supplementary table 1). For this, based on all the possible sub-datasets that fulfilled the specified criteria ensuring dataset integrity, we observed a relatively smooth increase in variability due to technical sources that progressed with increased number of technical replicates, experiments, and persons. However, importantly, the cumulative variability was almost doubled when data from two laboratories were combined. Adding a third laboratory to the dataset did not substantially increase the cumulative variability (Fig. 2c-d, Supplementary figure 5).

### Batch effect removal

Inspired by the extensive research in RNA-seq experimental designs to measure and correct for batch effects, we applied a similar approach to our study to curate the variability. For this, the LME model was computed using the complete dataset (both control and ROCK inhibition conditions), keeping the same random effects as previously used and including the control or ROCK inhibition as fixed effect.

We conducted this approach to the Instantaneous Cell Speed (ICS, Fig 3a-b) and to the first and second Principal Components of all variables (Fig 3c-d). For each observation, we computed and discriminated the effects derived either from random effects (derived from the lab, person, experiment, or technical replicate) or from the fixed effect (ROCK inhibition)^19^. The results clearly show that this approach allows for an unambiguous discrimination between the control and treatment conditions, therefore showing that the experimental variability in cell migration experiments can be addressed in order to better discriminate the effect of a given perturbation (Fig. 3a-d, supplementary figure 6). The batch-effect-removed data showed a robust increase in ICS as a result of the perturbation in the data from all three laboratories. In comparison, only laboratory #1 produced a similar sized increase as without batch effect removal, while the other labs displayed a small decrease (laboratory #2) or a small increase (laboratory #3). Thus, the direct comparison of data from cell migration experiments among our laboratories, each highly experienced in cell migration designs and experiments, could lead to discordant conclusions on the perturbation effect. This highlights the importance of statistical methods for batch effect removal in image-based quantitative studies.

**Figure 3.**
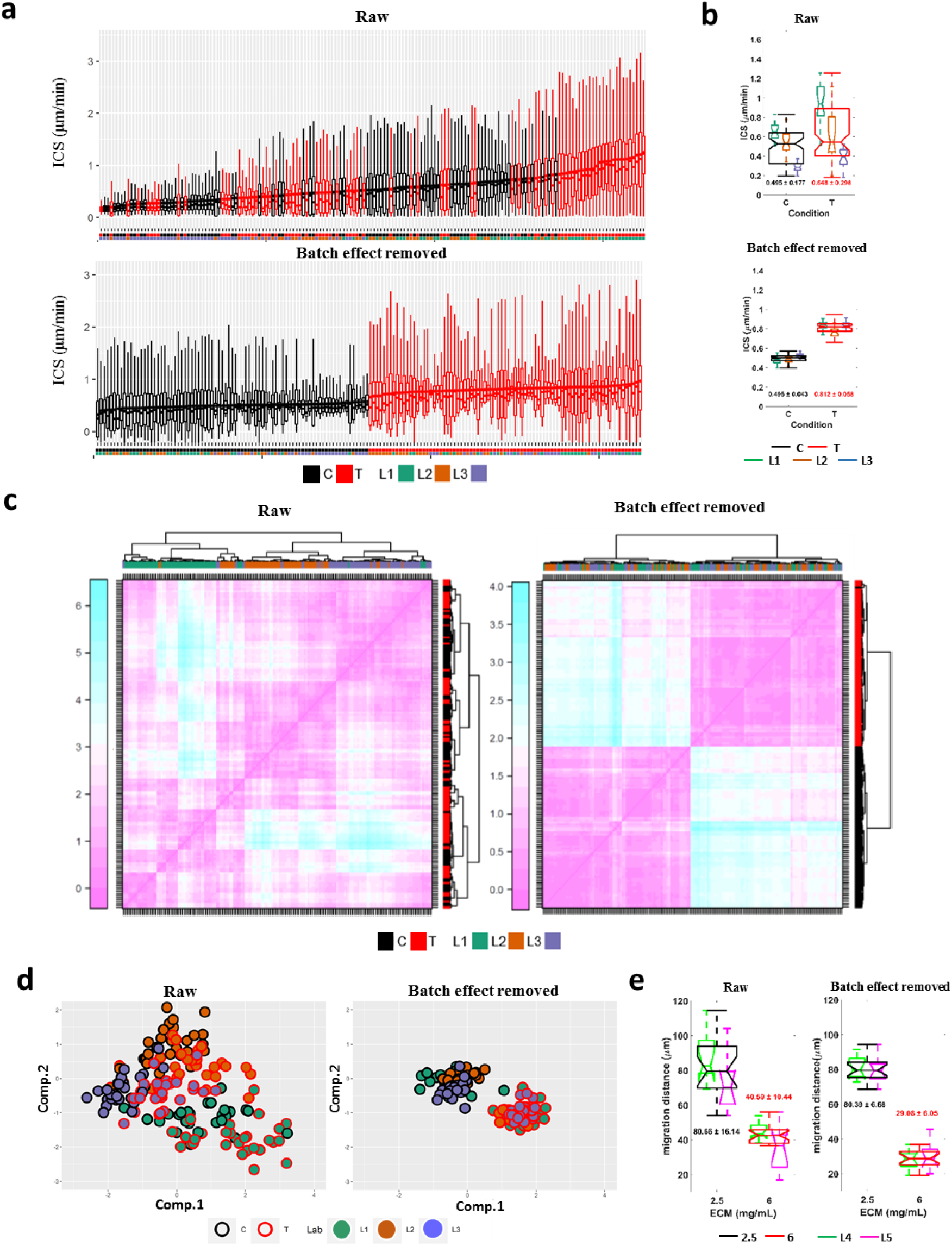
Batch effect removal dramatically reduces the variance. **a. Instantaneous Cell Speed (ICS) distribution before (top) and after (bottom) batch effect removal on control (C - black) and perturbed (ROCK inhibition) (T - red)**. Boxplots display ICS observations for each replicate, sorted by increasing value of the mean. Control and perturbation conditions are shown in black and red respectively. Laboratories in which each replicate was performed are color coded below the boxplots. **b. Mean ICS values and variance before (top) and after (bottom) batch effect removal**. Boxplot of mean ICS of each technical replicate from control and perturbed conditions in different laboratories. Laboratories are colour coded, while the aggregate results from all labs are shown in black (control) and red (perturbed). The numbers below the corresponding boxplot show mean ± standard division of the aggregated control/treated results from all labs. **c. Heatmap of the distance matrix before and after batch effect removal**. The heatmaps show average values of the distance matrix between 1st and 2nd Principal Components per lab, person, experiment, condition, and technical replicate before (left) and after (right) batch effect removal. Each row/column corresponds to one technical replicate. Sorting based on hierarchical clustering. **d. Batch effect removal in principal component data of 2D cell migration data**. Technical replicate of 1st and 2nd Principal Component average values before (left) and after (right) batch effect removal are shown in the same PCA space. Each dot represents one technical replicate. Results from different laboratories/conditions are colour coded as indicated. **e. Batch effect removal in 3D cell migration data**. Boxplot shows the mean 3D cell migration distance of HT1080 cells embedded in different concentrations of collagen before (left) and after (right) batch effect removal. Different ECM concentrations are shown in black (2.5 mg/mL) or red (6 mg/mL) and data from different laboratories are indicated with green (Laboratory #4) and magenta (Laboratory #5). The sums of results from both labs are shown in black (2.5 mg/mL) and red (6 mg/mL). The mean ± standard division of the aggregated 2.5mg/mL or 6 mg/mL results from both labs are shown with the corresponding boxplot. For the boxplots in **a, b**, and **e**, in each box, the central mark indicates the median, and the bottom and top edges of the box indicate the 1^st^ quartile and 3^rd^ quartile respectively.

We also applied the batch effect removal approach to a 3D cell migration dataset generated from two independent laboratories with a similar strategy as for the 2D cell migration experiment (supplementary figure 7). The difference of migration distance of the cells in response to low (2.5 mg/ml) or high density (6 mg/ml) concentration of polymerized collagen was already reliably discriminated comparing the raw data (supplementary figure 8), as previously described ^20^. However, significant lab-to-lab variance of results within each test group was still observed (Fig. 3e left). Also in this case, the batch effect removal processing significantly reduced the variance and provided a more robust difference (Fig. 3e right).

## Discussion

The emerging increase in high content imaging data sharing provides opportunities for data reuse and meta-analysis, but the usefulness of these opportunities remains largely untested, and the sources of variance within this type of data have not been characterized. In this study, we found that variation between laboratories is the major sources of technical variance in high content imaging data of cell morphology and migration features. This outcome suggests that, although the experimental design was idealized including sharing of a detailed protocol, cells and reagents, standardizing details such as cell passaging prior to the experiment, cell density prior to seeding for migration, the type of fetal bovine serum, and cumulative passage number of cells, the lab-to-lab variance currently limits the value of meta-analysis of the basic high content cell image data. This lab-to-lab variance may at least in part be explained by observed local variations in equipment and practices, including use of different microscopes and their differences in what imaging plates could be harbored, and lab-to-lab differences of cell density apparent in the images, despite that the same standard method was used for cell quantification.

Importantly however, we show that application of a batch effect removal approach significantly reduced the technical variance at all levels and provided useful meta-analysis of perturbation effects in both 2D and 3D spheroid culture models performed in different laboratories, at least under our highly standardized conditions. Similar batch effect removal approaches have been important for meta-analysis in other fields and data types, such as from RNA-sequencing and peptide-centered proteomics via mass spectrometry^21-23^.

Taken together, our study indicates that the usefulness of high content image data meta-analysis is currently limited to the study of perturbation effects, and for which batch effect removal is necessary. Our study entailed a high degree of standardization. Further studies are therefore needed to define the usefulness of meta-analysis of more typical high content image datasets that are more loosely standardized than ours and that often differ not only in their precise design, but also in study purpose and aim.

## Methods

### Cell culture and imaging

#### 2D cell migration

We developed highly detailed protocols for cell culture and seeding for live cell imaging that was shared and used for all experiments (see Supplementary material 2). Briefly, the Friedl laboratory provided HT1080 cells stably expressing LifeAct-mCherry & H2B-EGFP (before sharing, the Sahai laboratory performed the standard cell authentication procedure on this cell line by comparing its STR profile to the public database). Mycoplasma infection was excluded prior to the experiments. Cells were cultured with high glucose DMEM supplemented with FBS (10%), sodium pyruvate (1mM) and penicillin/streptomycin (100 U/ml). Cells were passaged at ∼80 - 90 % confluence, up to passage number 20. One day before imaging, 2 × 10^5^ cells were seeded onto one well of the 6 well plates and left overnight in the incubator. On the experimental day, the assay wells were prepared as follows: 100 μl of 20 μg/ml collagen I was added to each of six wells in a 96-well imaging plate or chambered coverslip and incubated at 37°C for 2 h. The supernatant was discarded by flipping the plate upside down and subsequently, each well was incubated at 37°C for 20 min with 100 μl of heat denatured 0.5% BSA for blocking. 500 cells in 100 μl of serum free culture medium were seeded in each of the six individual wells of the 96-well imaging plate or chambered coverslip, ensuring homogeneous cell distribution by tapping the plate or chambered coverslips in perpendicular directions. After 10 min, during which cells attached to the well bottom, the imaging plate was incubated at 37°C and 5% CO_2_ for 2.5 h.

For the live cell imaging, we used multidimensional automatized microscopes with an environmental chamber to keep temperature, humidity, and CO_2_ constant. Pre-warmed media with or without ROCK inhibitor (Y27632, final concentration at 15 μM) was added before the start of imaging. A 20x 0.75 NA objective was used and tiled images (5 × 5) were generated to capture a large area in each well. The images were acquired in 5 min interval for 6 h.

The detailed protocol is attached in the supplementary material (Supplementary Material 2-4). Any deviations from the distributed procedure were recorded and summarized (Supplementary Material 5).

#### 3D spheroid invasion assay

We developed a detailed workflow for a 3D spheroid invasion assay that was shared and used for all experiments performed at three different locations. For detailed protocols for 3D spheroid culture and labeling, imaging, and image analysis, see Supplementary Material 6, 7, and 8, respectively.

##### 3D spheroid culture and labeling

Briefly, the Friedl laboratory provided HT1080 cells. Before sharing among the three groups, the Sahai laboratory validated this cell line by comparing its STR profile to the published ones. Mycoplasma infection was excluded prior to the experiments. Cells were cultured in T75 flask with 10% CO_2_ at 37°C. Multicellular spheroids containing 1000 HT1080 cells were generated using hanging-drop culture method^24. 24. 24^.The spheroids were embedded in rat tail collagen I (Corning, Cat no. 354249), in up to 18wells of 96-well imaging plates per collagen concentration, using 1 spheroid per gel and a final collagen concentration of 2.5 or 6 mg/ml. Former protocols for spheroid embedding^20, 25^ were adapted to have control over the number of spheroids per well, spheroid height with respect to imaging window and the onset of collagen polymerization, to minimize variation between technical repeats per plate. Plates were incubated for 24 hours at 37°C to establish cancer cell invasion in three dimensions, prior to fixation in 4 % PFA. The 3D cell cultures were fluorescently stained with DAPI (Sigma, D9542, 2 μg/ml) and AlexaFluor633-Phalloidin (Molecular Probes, A22284, 1:200 dilution) and stored (preferably for <48 hours) at 4°C prior to imaging.

##### Imaging

In brief, the lower left corner of the spheroid was positioned in the scan field, with the border of the spheroid core touching the image border. A z-range of up to 120 μm was used to image from z = 1/2 to z = 4/5 of spheroid dimensions. Transmission, reflection and fluorescence channels were recorded sequentially at 8-bit resolution. The laser power was set close to the saturation limit of the dye. The detector amplification (high voltage) was set in such a manner, that the brightest cells in migration zones made use of the full digital detection range. In both laboratories, imaging was performed using a Zeiss LSM880 equipped with a 20x 0.8 NA objective. The following microscope parameters were used: scan field 708.5μm^2^, pixel size 1.2 μm, pixel dwell time 1.3 μs, z-step size 2 μm and line averaging 3.

All the metadata of the images were also recorded.

### Image processing

#### 2D live cell imaging data

Images were converted from the original format to.tif format. To generate large image composites, stitching was performed either automatically during acquisition or via a custom-made MatLab script. Images from different laboratories were resized to the same resolution (0.8260 μm/pixel), to allow proper further comparison. A CellProfiler (v 2.2.0) pipeline was used to automatically segment and track cells and nuclei, and to extract 15 morphological and dynamic variables from the raw images (shown in Figure 2a). Because images from different laboratories were acquired with different types of microscopes, threshold correction factors for the segmentation on cells and nuclei were adapted to the data from different labs.

In order to identify protrusions, retractions, and short-lived cell regions, we compared consecutive, segmented cell images from the CellProfiler analysis results using tailored Matlab scripts. Protrusions were identified as regions present in a cell at a certain time point but absent in the previous. Retractions were defined as regions present at one time point but absent in the next time point. Short-lived regions are those regions that are present at only one time point but not in the ones directly before or after, corresponding to a lifetime of <10 min ^17^.

The CellProfiler pipelines for each laboratory and the Matlab scripts are available together with the raw images at the SciLifeLab Data Repository (https://doi.org/10.17044/scilifelab.21407402).

#### spheroid invasion data

This workflow is implemented in Fiji as the Nucleus Annotation 3D (NA) and the Cell Migration Analyser 3D (CMA) plugin sets (https://github.com/Mverp/Nucleus-Annotation-3D and https://github.com/Mverp/Cell3DMeasurements) and was distributed to 2 independent labs (RUMC and CRICK) for standardized analysis of independent datasets from spheroid culture performed in each lab independently (Supplementary material 8). First, the outline of the spheroid core was defined by manually setting four points far away from each other, in the 3D image stack to be analyzed, at the spheroid border. Based on the four points, the annotation program defined a sphere in the dataset, which was used as a reference for migration distance from the spheroid core. Then, the DAPI channel of the 3D image datasets was used for nuclear segmentation, segments at the border were removed and the distance of the center of each nucleus to the defined spheroid core was quantified and recorded for subsequent analysis.

To optimize and validate the plugin, segmentation outputs were compared to manually annotated ‘gold standard’ images. After segmentation, the nuclei occurring in both annotation and segmentation output, the true positives, were automatically calculated. Next, the performance of the segmentation was quantified by calculating the precision (# true positives / # nuclei in segmentation) and recall (# true positives / # nuclei in the annotation). The analysis was performed on the image data of both labs using the optimized settings (MigrationAnalysisParameters.txt, included in the SciLifeLab Data Repository https://doi.org/10.17044/scilifelab.21407402).

### Data processing and modelling of the 2D cell migration data

Pre-processing: Based on the original tracking data, the static and rounded cells were excluded based on visual assessment. Then the duplicated and merged cell/nuclear trajectories were identified and removed. Excessively large (cell area > third quantile + 1.5*inter quantile range of the areas from all cells) or small (nuclear area < 100 μm^2^) cells were excluded based on the measurements of cellular and nuclear area, in order to remove noise from cell debris and cell aggregations. The remaining trajectories were smoothed with the rolling window method with window size of 9. The Instantaneous Cell Speed (ICS) was calculated based on the smoothed trajectories.

The linear mixed effect modelling was performed based on the R package *lme4*^*26*^.

For the cumulative variability calculation, we designed an hypothetical experimental design with increasing levels of complexity: 2 or 3 replicates, 2 or 3 experiments with 3 replicates each, 2 or 3 persons performing 3 experiments with 3 replicates each, or 2 or 3 labs where 3 persons perform 3 experiments with 3 replicates. For this, we generated all the possible sub-datasets which fulfilled the specified criteria ensuring dataset consistency (this is, we avoided the combination of data which did not have the same origin in the higher level of the hierarchical structure), and computed the cumulative variability for each level. Supplementary table 1 shows how the datasets were generated. As a control, we firstly randomized the original data and then generated similar sub-datasets as the original ones and calculated the cumulative variability in the same way.

### Batch effect removal

After fitting the original data with linear mixed effect model to extract the fixed and random effects, each single observation was modified by subtracting the intercept from all levels (lab, person, experiment, technical replicate, and observation) and adding the fixed effect between two conditions (control vs perturbation in 2D migration; 2.5 mg/mL vs 6 mg/mL collagen in 3D invasion).

### Statistical analysis

Statistical analysis was performed using R (R Core Team).

## Supporting information

Protocol for HT1080 2-D Migration_Live Cell Imaging

Control Sample Movie

ROCK Inhibitor Sample Movie

2D Migration_survey summary

3D spheroid culture protocol

3D spheroid microscopy protocol

3D spheroid image analysis protocol

## Data availability

All of the raw images, survey, image analysis pipelines, and data analysis scripts are shared in the SciLifeLab Data Repository (https://doi.org/10.17044/scilifelab.21407402).

## Acknowledgements

Acknowledgement to funding from the European Union’s Horizon 2020 Programme under the MultiMot project, Grant Agreement 634107 (PHC32–2014), and to SS from the Swedish Research Council, The Strategic Research Foundation (Sweden), and the Swedish Cancer Society. The authors thank Vito Conte for the help in scientific notation for cumulative variability analysis. The authors from Karolinska Institutet, Stockholm (KI) performed imaging at the Live Cell Imaging core facility/Nikon Center of Excellence, KI, Sweden supported by grants from the KI infrastructure committee. The Ghent University, Gent (UGENT) authors thank the VIB BioImaging Core for support and access to the instrument park. The Radboud University Medical Centre, Nijmegen (RUMC) authors thank Esther Smeets for her contribution to the 3D image analysis pipeline and, Jeroen Slaats and Jan-Hendrik Veenhuizen for manual annotation of the 3D image data. Furthermore, they thank the Research Technology Centre for Microscopy at the RUMC for support and access to the confocal microscopes.

## Contributions

Standardization development and execution was performed by the laboratories at KI, RUMC, the Sir Francis Crick Institute, London (CRICK), UGENT and the Weizmann Institute of Science, Rehovot (WI). For the 2D experiment, KI established and shared the protocol, materials and reagents, and coordinated data collection and analysis. For the 3D experiment, RUMC coordinated the development of the protocol, the experimental material sharing, and the data collection. RUMC, UGENT, and CRICK established and shared the protocol, materials and reagents, and data analysis. RUMC provided HT1080 cells stably expressing LifeAct-mCherry & H2B-EGFP.

SS conceived and supervised the overall study.

JH designed and distributed the 2D cell migration experiment protocol and materials.

PF designed the 3D cell migration experiment. GJB coordinated the 3D cell migration experiments. PF, GJB, MVT, NE, MV and JC developed and optimized the 3D hanging drop spheroid assay further for high-content imaging.

JH, XG, YD, SWK, IG, OP, MVT, KB, EVH, GJB, MV, NE, JC, and AMD performed the experiments.

XSP designed and implemented the image processing and quantification for the 2D experiment.

MVE and GJB designed the image analysis protocol for the 3D experiment. MVE programmed the scripts and FIJI plugins for the 3D image analysis. MVE, MVT, JC, and GJB applied the image analysis scripts onto the 3D image data. FW reviewed and corrected the established 3D experiment and image analysis protocols.

XSP designed and implemented the data and statistical analysis. JH implemented data and statistical analysis. MB supervised the statistical analysis and modelling.

JH, XSP and SS drafted the manuscript.

All authors read and approved the final manuscript.

