## Supplementary Figures and Supplementary Table for "Multi-site assessment of reproducibility in high-content live cell imaging data"

**Supplementary figures and tables**

**Supplementary Fig. 1**

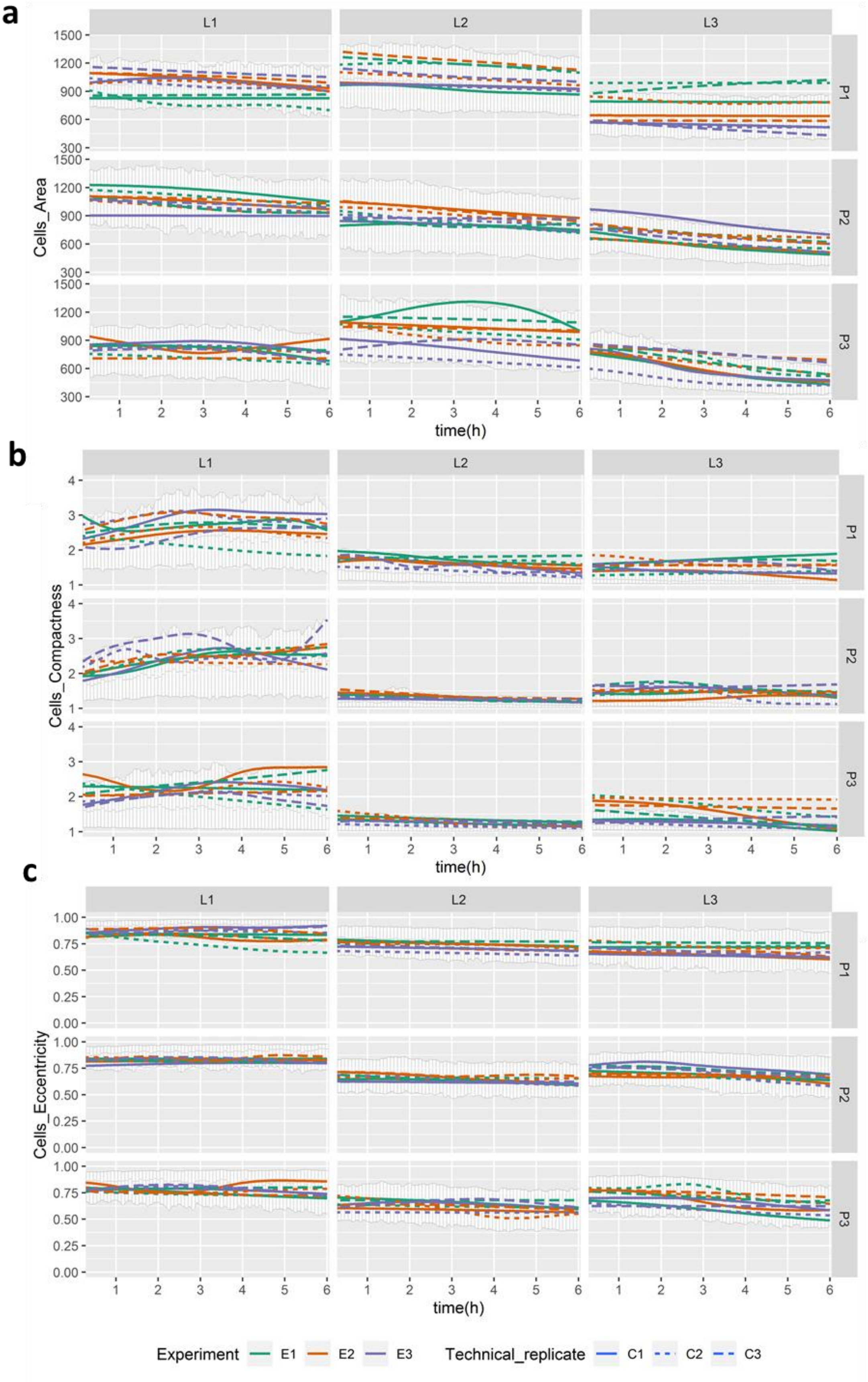

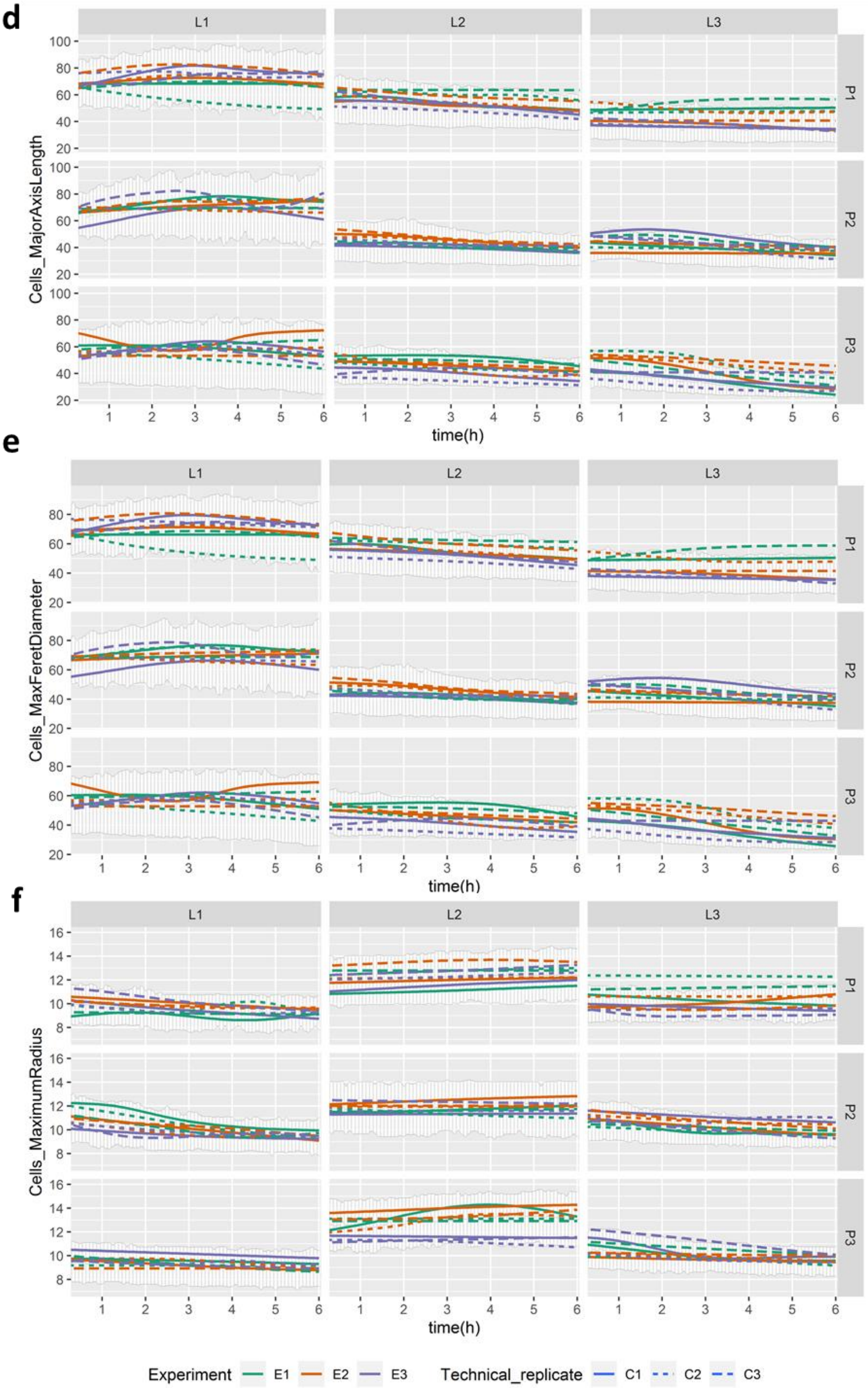

d
e
f
L1
L2
L3
P1
P2
P3
Cells_MajorAxisLength
Cells_MaxFeretDiameter
Cells_MaximumRadius
time(h)
Experiment
E1
E2
E3
Technical_replicate
C1
C2
C3

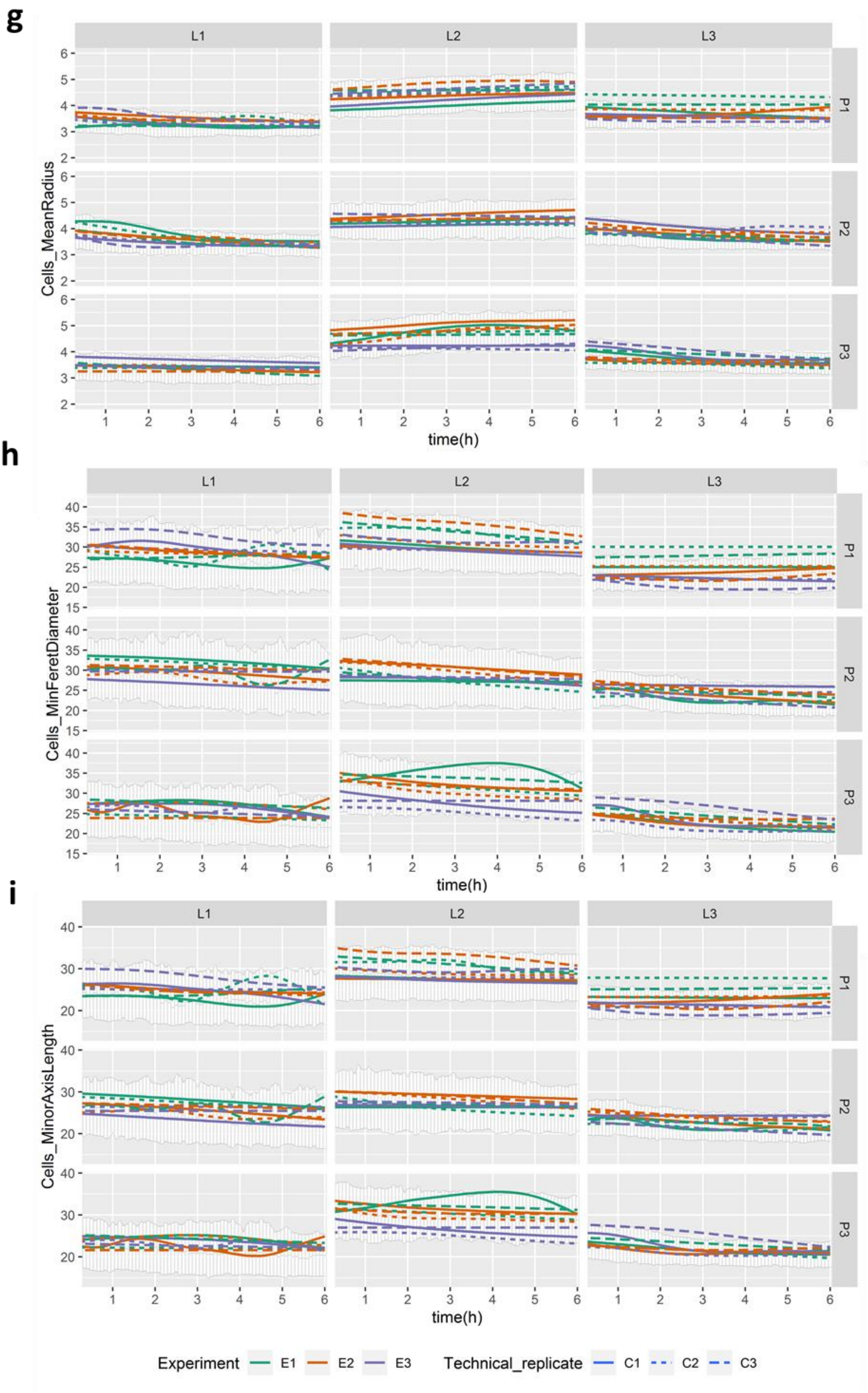

g
h
i
L1
L2
L3
P1
P2
P3
Cells_MeanRadius
Cells_MinFeretDiameter
Cells_MinorAxisLength
time(h)
Experiment
E1
E2
E3
Technical_replicate
C1
C2
C3

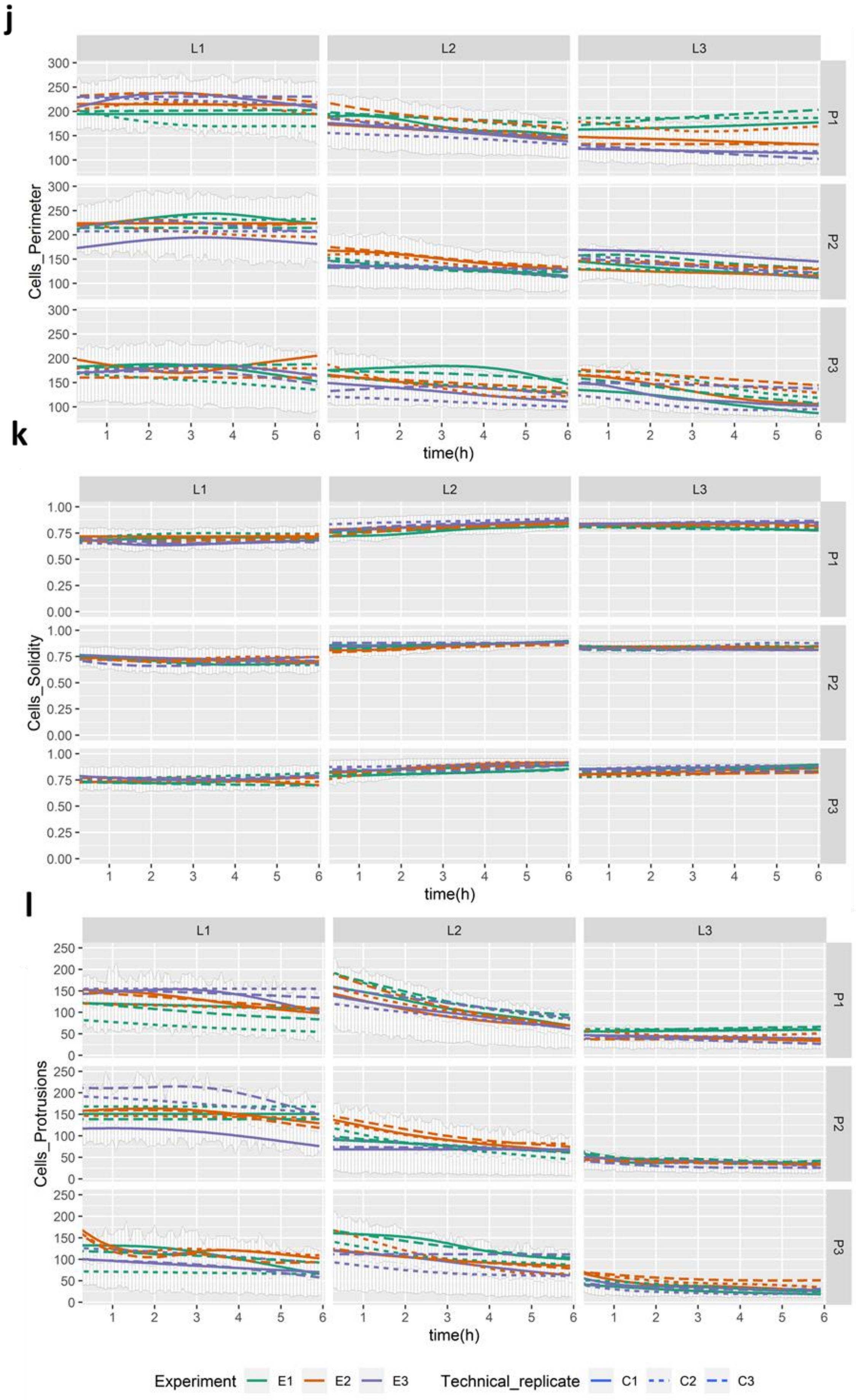
j
k
l
L1
L2
L3
P1
P2
P3
Cells_Perimeter
Cells_Solidity
Cells_Protrusions
time(h)
Experiment
E1
E2
E3
Technical_replicate
C1
C2
C3

**m**

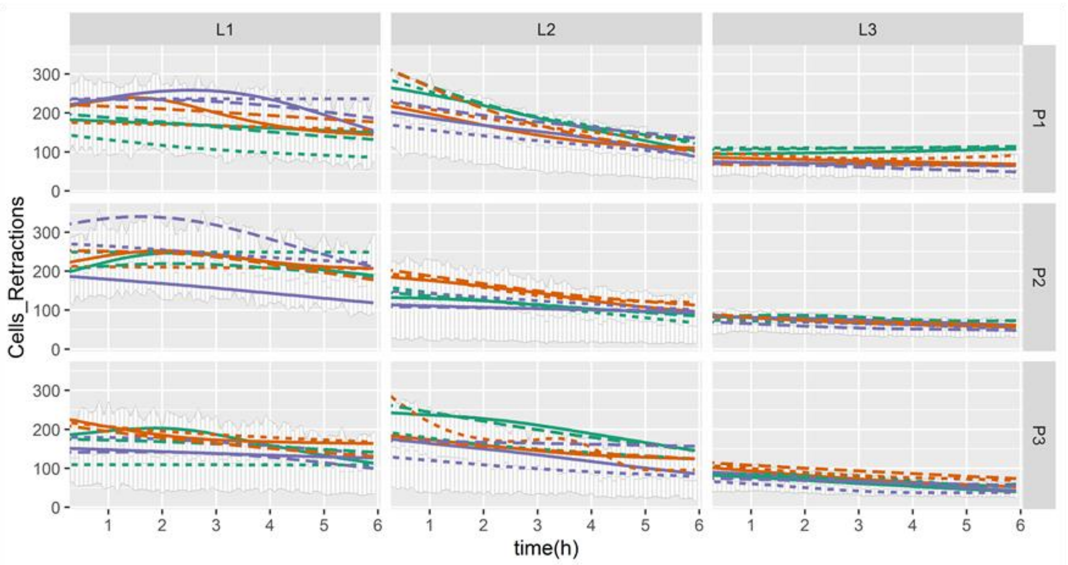
L1
L2
L3
P1
P2
P3
Cells_Retractions
0
100
200
300
1
2
3
4
5
6
time(h)

**n**

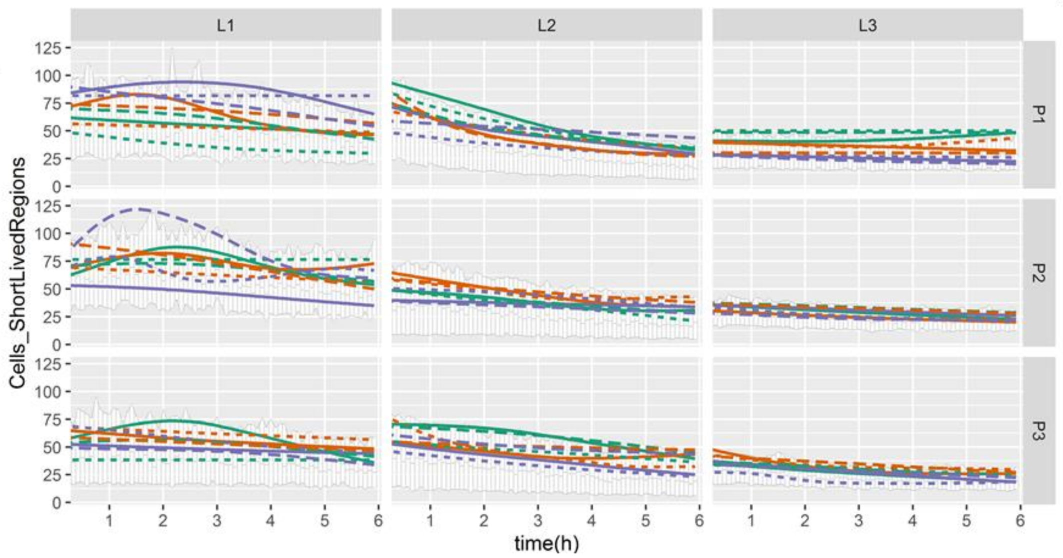
L1
L2
L3
P1
P2
P3
Cells_ShortLivedRegions
0
25
50
75
100
125
1
2
3
4
5
6
time(h)

**o**

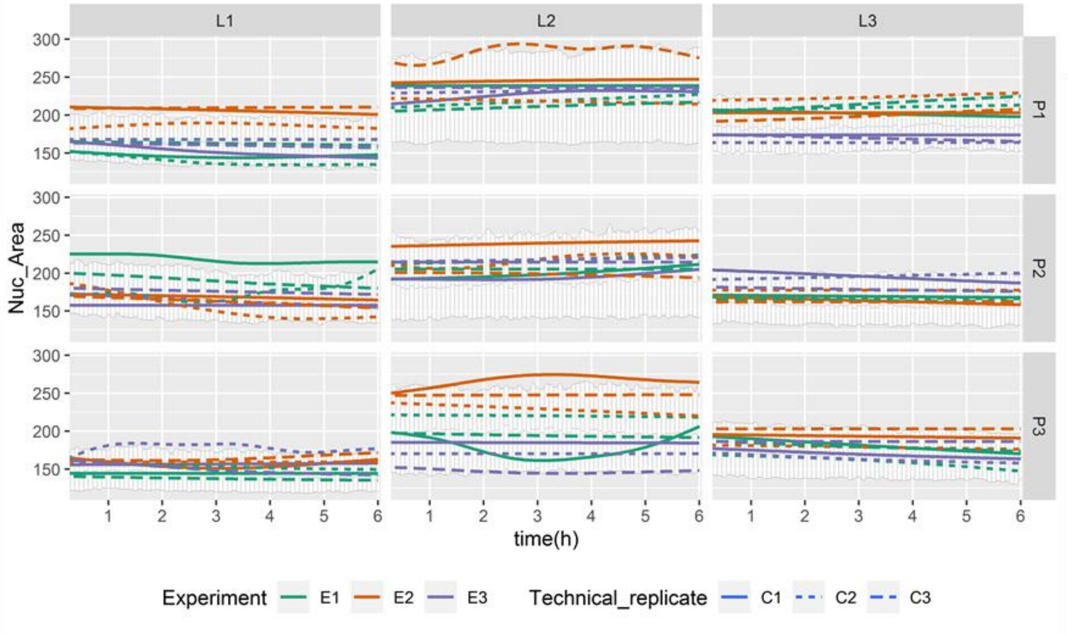
L1
L2
L3
P1
P2
P3
Nuc_Area
150
200
250
300
1
2
3
4
5
6
time(h)
Experiment E1 E2 E3
Technical_replicate C1 C2 C3

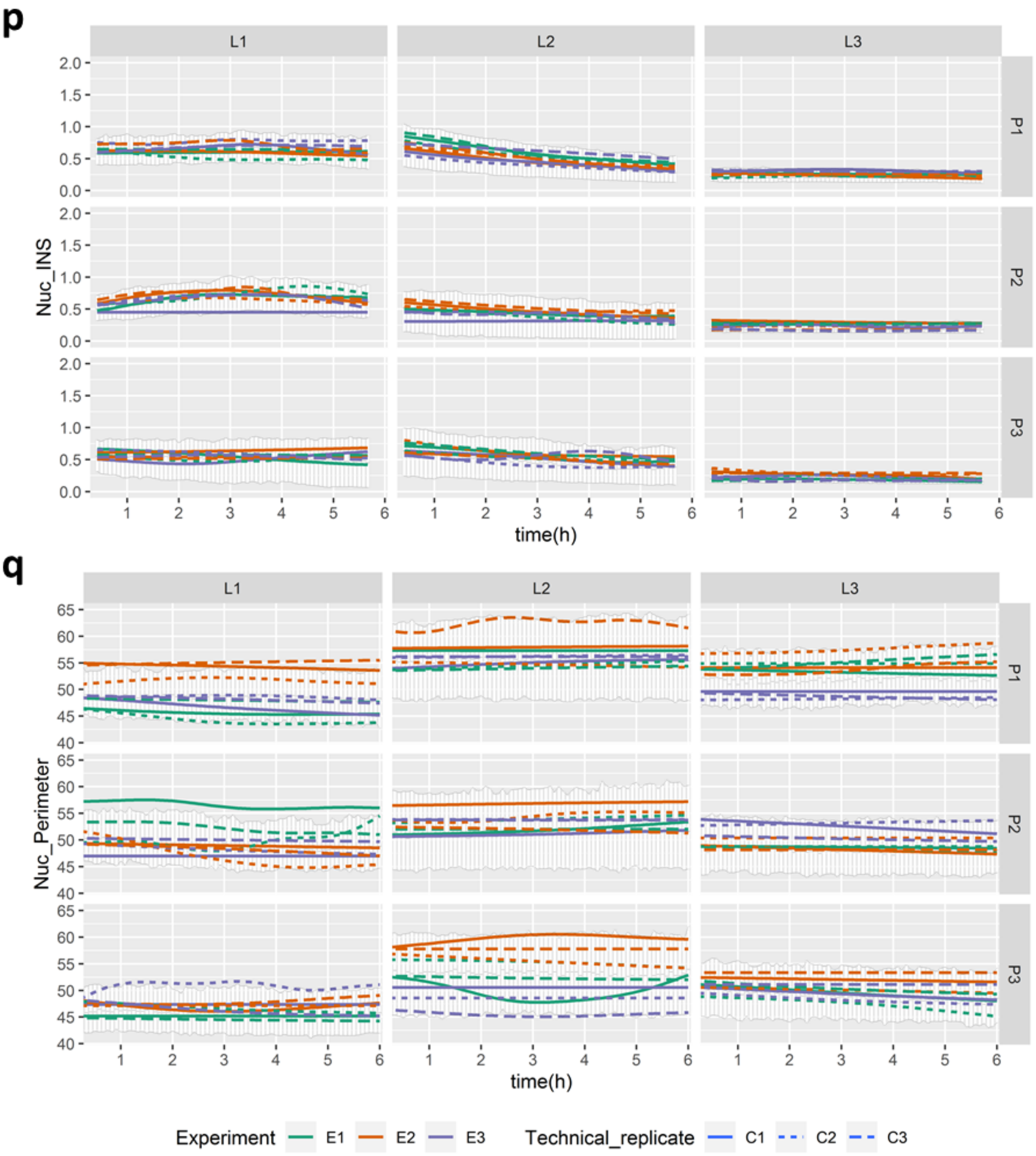


**Supplementary Figure 1. Quantification of variables over time at different levels of data hierarchy.** The graphs display quantifications over time for each laboratory (L1-3), person (P1-3), experiment (E1-3), and technical replicate (C1-3), of the control condition. Results include Cell Area(**a**), Cell Compactness(**b**), Cell Eccentricity(**c**), Cell Major Axis Length(**d**), Cell Maximal Feret Diameter(**e**), Cell Maximum Radius(**f**), Cell Mean Radius(**g**), Cell Minimal Feret Diameter(**h**), Cell Minor Axis Length(**i**), Cell Perimeter(**j**), Cell Solidity(**k**), Cell Protrusions(**l**), Cell Retractions(**m**), Cell Short Lived Regions(**n**), Nuclear Area(**o**), Nuclear Instantaneous Nuclear Speed (INS) (**p**), and Nuclear Perimeter(**q**). Lines in different colours represent three different experiments. Different style of the lines with the same colour represent three different technical replicates from the control condition within one experiment. The error bar indicates the first and third quartiles of the data at each time point.

**Supplementary Fig. 2**

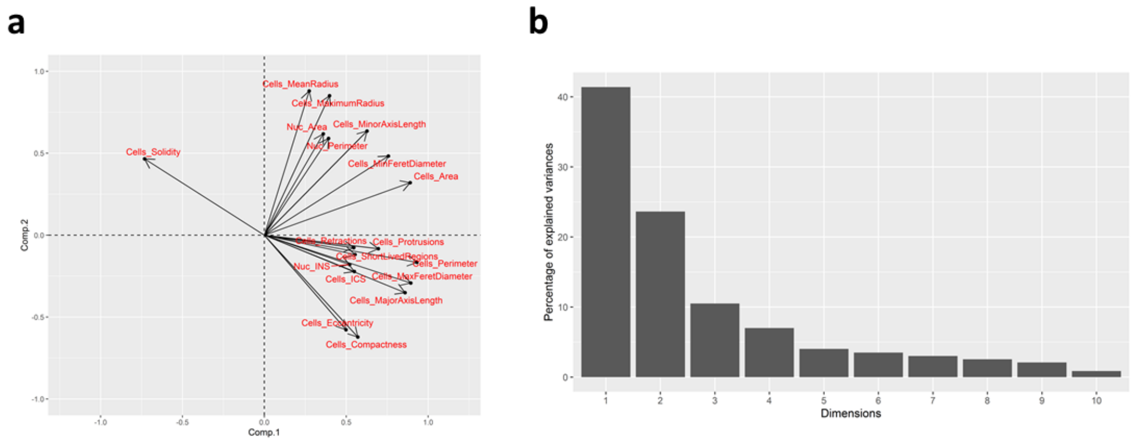


**Supplementary Figure 2. Overview of the Principal Component Analysis (PCA) results from the 18 variables used in the study. a**. **Variance map of the PCA analysis.** Each arrow shows the relative location of the corresponding variable within the principal component space. **b**. **Percentage of explained variance by the top 10 principal components**. X axis shows the dimension of the principal component. Y axis shows the percentage of the explained variances by the corresponding principal component.

**Sup Fig. 3**

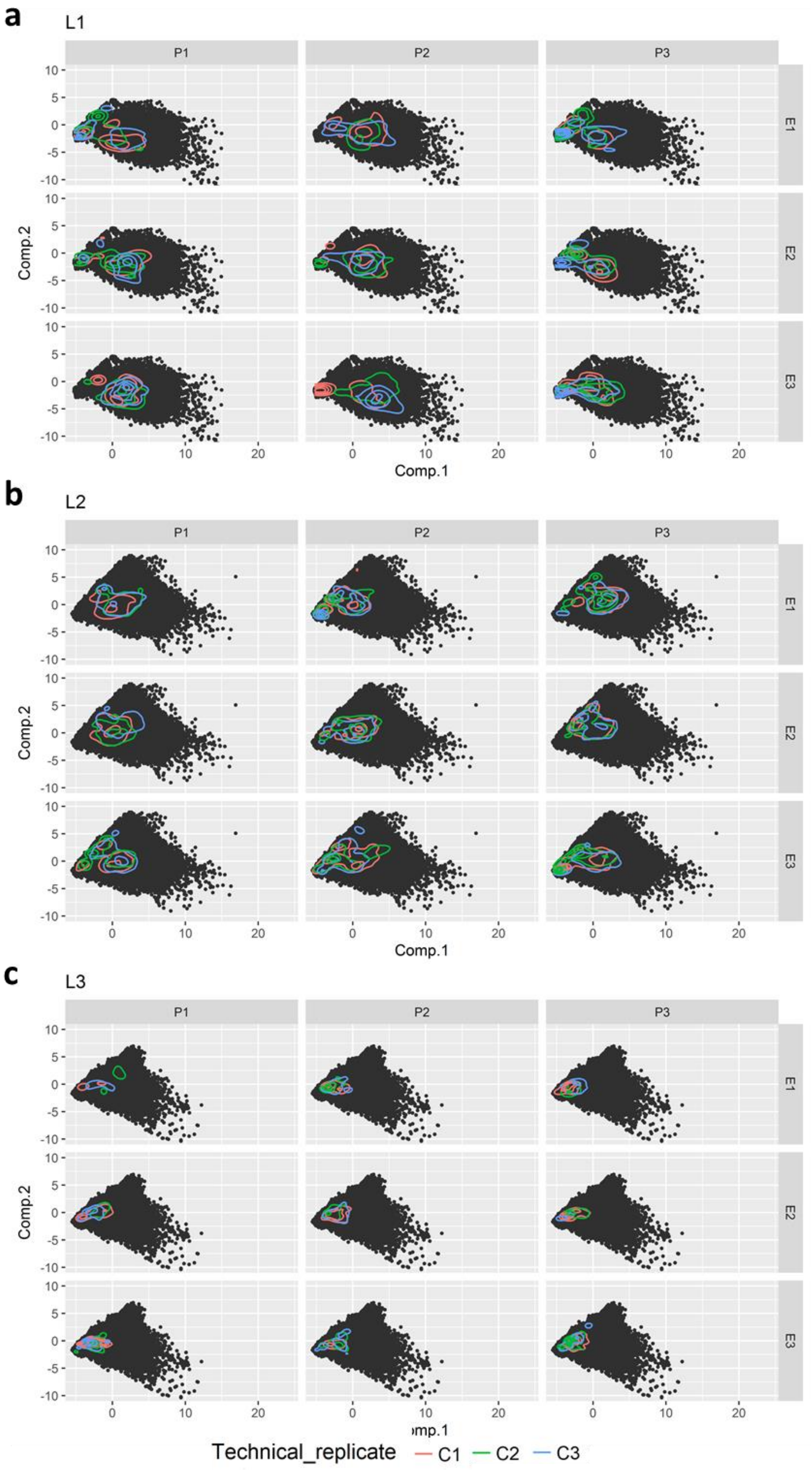

a
L1
b
L2
c
L3
P1
P2
P3
E1
E2
E3
Comp.1
Comp.2
Technical_replicate
C1
C2
C3

**Supplementary Figure 3. Principal component analysis results shown for each experiment from each person in three laboratories.** Principal component analysis in individual experiments of each person in Laboratory #1(**a**), Laboratory #2(**b**), and Laboratory #3(**c**). Black dots show the position of the first and second principle components for each observation. Coloured lines show the 2D density plots of the technical replicates. Each technical replicate within the same experiment is shown in different colours. The principal component space is identical for all the plots.

**Sup Fig. 4**

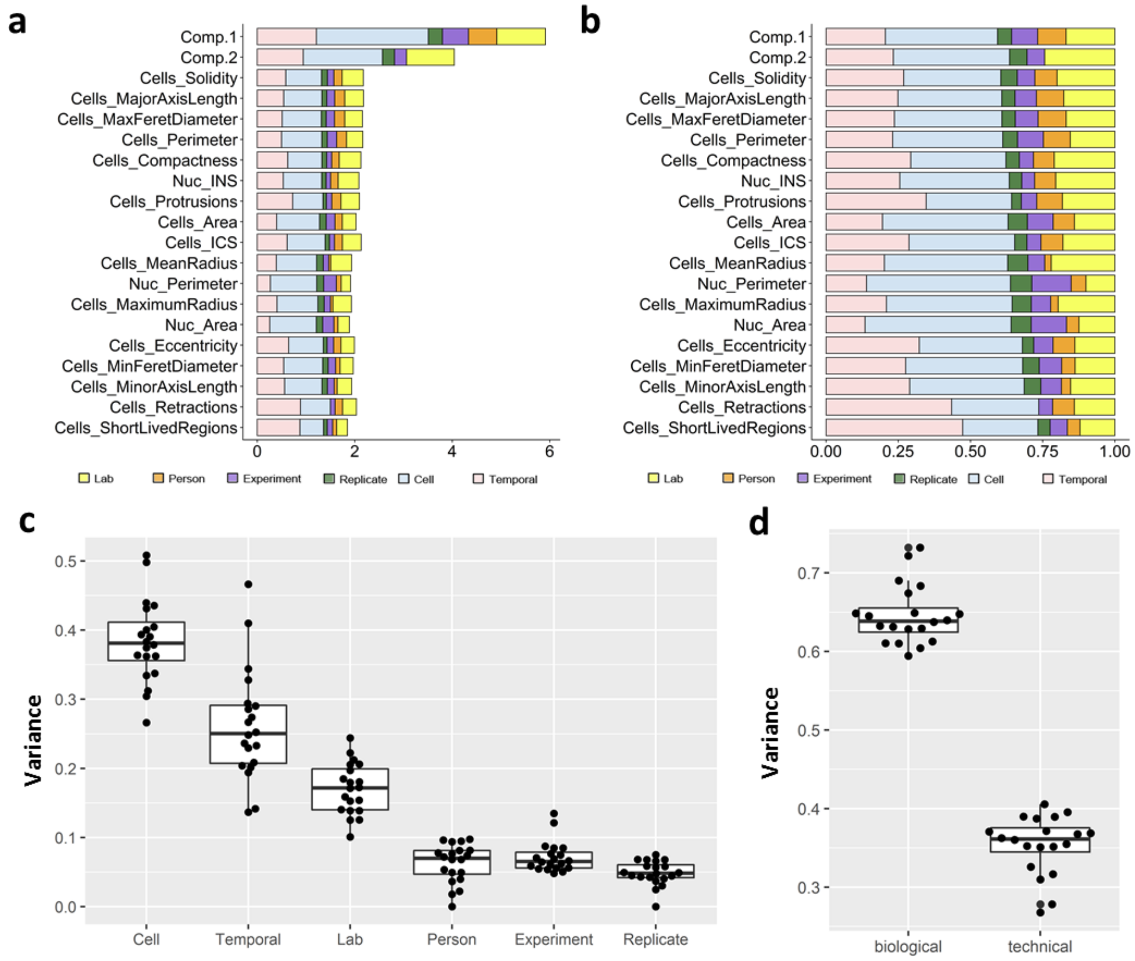


**Supplementary Figure 4. Variance components of each variable from all data from the control condition hierarchical levels based on the Linear Mixed Effect (LME) model analysis. a-b.** Absolute (**a**) or relative (**b**) variance components of each variable from temporal, cell, technical replicate, experiment, person, and laboratory levels based on the LME model analysis. **c**. Boxplot of the absolute variance components of all the variables from temporal, cell, technical replicate, experiment, person, and laboratory levels based on the LME model analysis. Each dot represents one variable within the corresponding variance level. **d**. Boxplot of the biological (cell and temporal) and technical (technical replicate, experiment, person, and lab) variance components of all the variables based on the LME model analysis. Each dot represents one variable within the corresponding variance category. For the boxplots in **c** and **d**, on each box, the central mark indicates the median, and the bottom and top edges of the box indicate the 1st quartile and 3rd quartile, respectively. The whiskers below and above the box indicate the 1st quartile-1.5* interquartile range, and 3rd quartile+ 1.5* interquartile range, respectively.

**Supplementary Fig. 5**

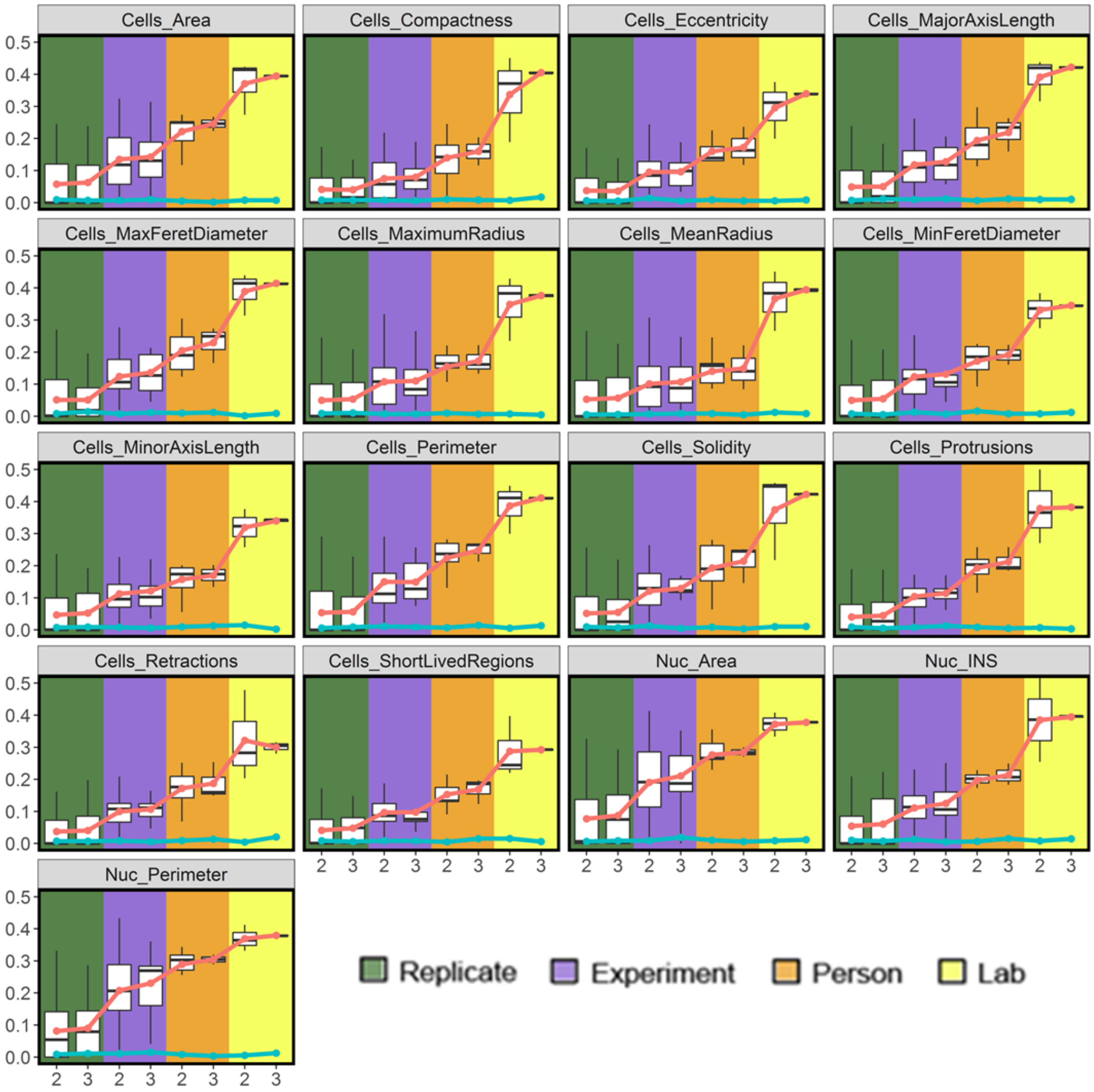


**Supplementary Figure 5. Cumulative variance of the different hierarchical data levels.** Each graph displays the cumulative variance of increasing data hierarchy, including technical replicate (R, green), experiment (E, purple), person (P, orange), and laboratory (L, yellow) levels. Results include Cell Area, Cell Compactness, Cell Eccentricity, Cell Major Axis Length, Cell Maximal Feret Diameter, Cell Maximum Radius, Cell Mean Radius, Cell Minimal Feret Diameter, Cell Minor Axis Length, Cell Perimeter, Cell Solidity, Cell Protrusions, Cell Retractions, Cell Short Lived Regions, Nuclear Area, Instantaneous Nuclear Speed (INS), and Nuclear Perimeter at technical replicate, experiment, person, and laboratory levels. Boxplots show variances with 2 or 3 replicates, experiments, persons, or laboratories, calculated at each level. Red dots show the mean value of the cumulative variance that are linked with red lines. As a control, cyan dots and lines show the cumulative variance of the same data after randomization. For the boxplots in each subfigure, on each box, the central mark indicates the median, and the bottom and top edges of the box indicate the 1$^{st}$ quartile and 3$^{rd}$ quartile respectively. The whiskers below and above the box indicate the 1$^{st}$ quartile-1.5* interquartile range, and 3$^{rd}$ quartile+ 1.5* interquartile range, respectively.

**Supplementary Fig. 6**

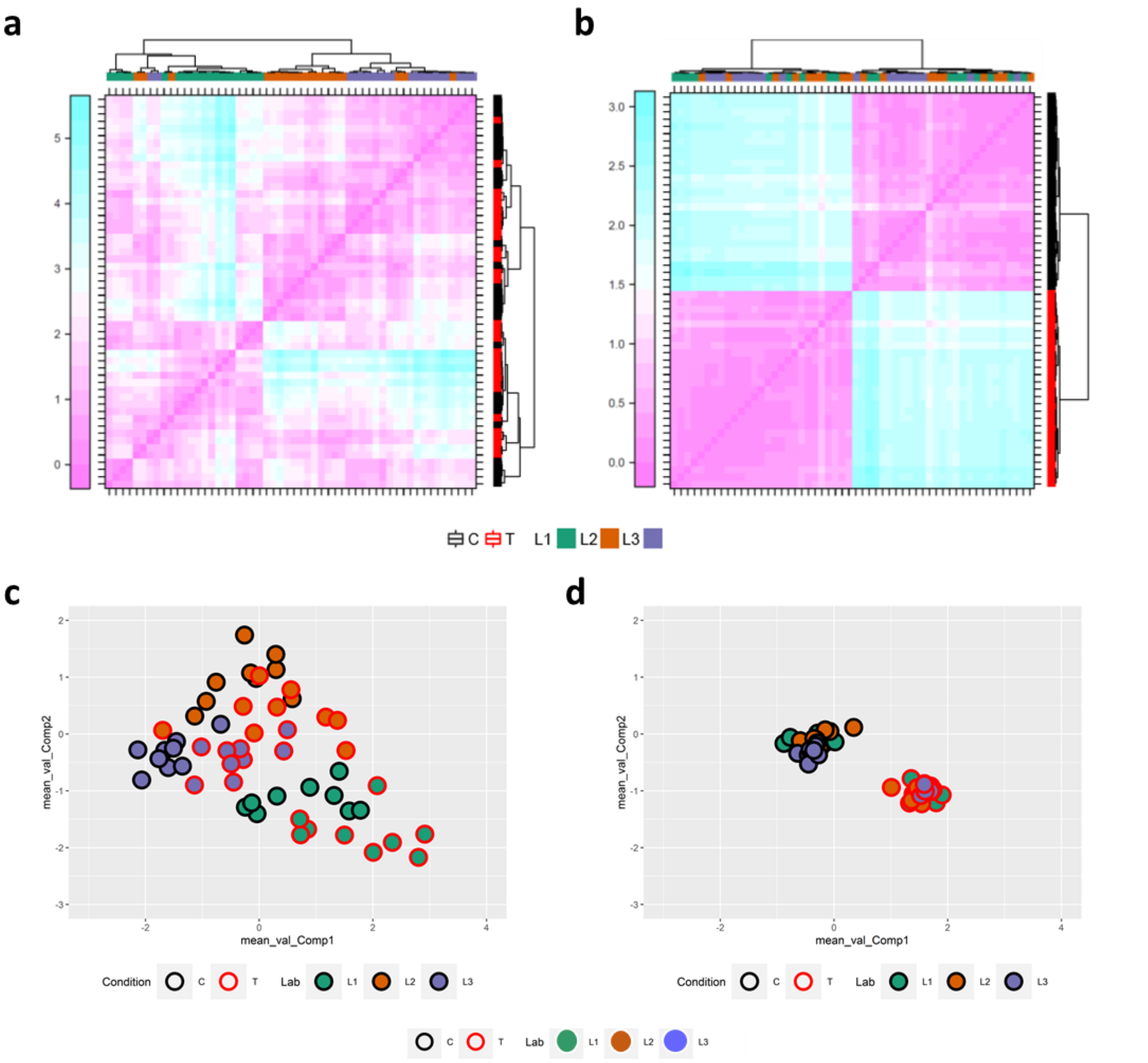


**Supplementary Figure 6**. **Heatmap of the distance matrix before and after batch effect removal. a-b** Average values of the distance matrix between 1st and 2nd Principal Components per lab, person, experiment, and condition before (**a**) and after (**b**) batch effect removal are shown in the heatmaps. Each row/column corresponds to one experiment. Sorting based on hierarchical clustering. **c-d**. Average values of 1st and 2nd Principal Components at the experiment level before (**c**) and after (**d**) batch effect removal. Each dot represents one experiment. Results from different labs/conditions are coded with different colors as indicated.

**Supplementary Fig. 7**

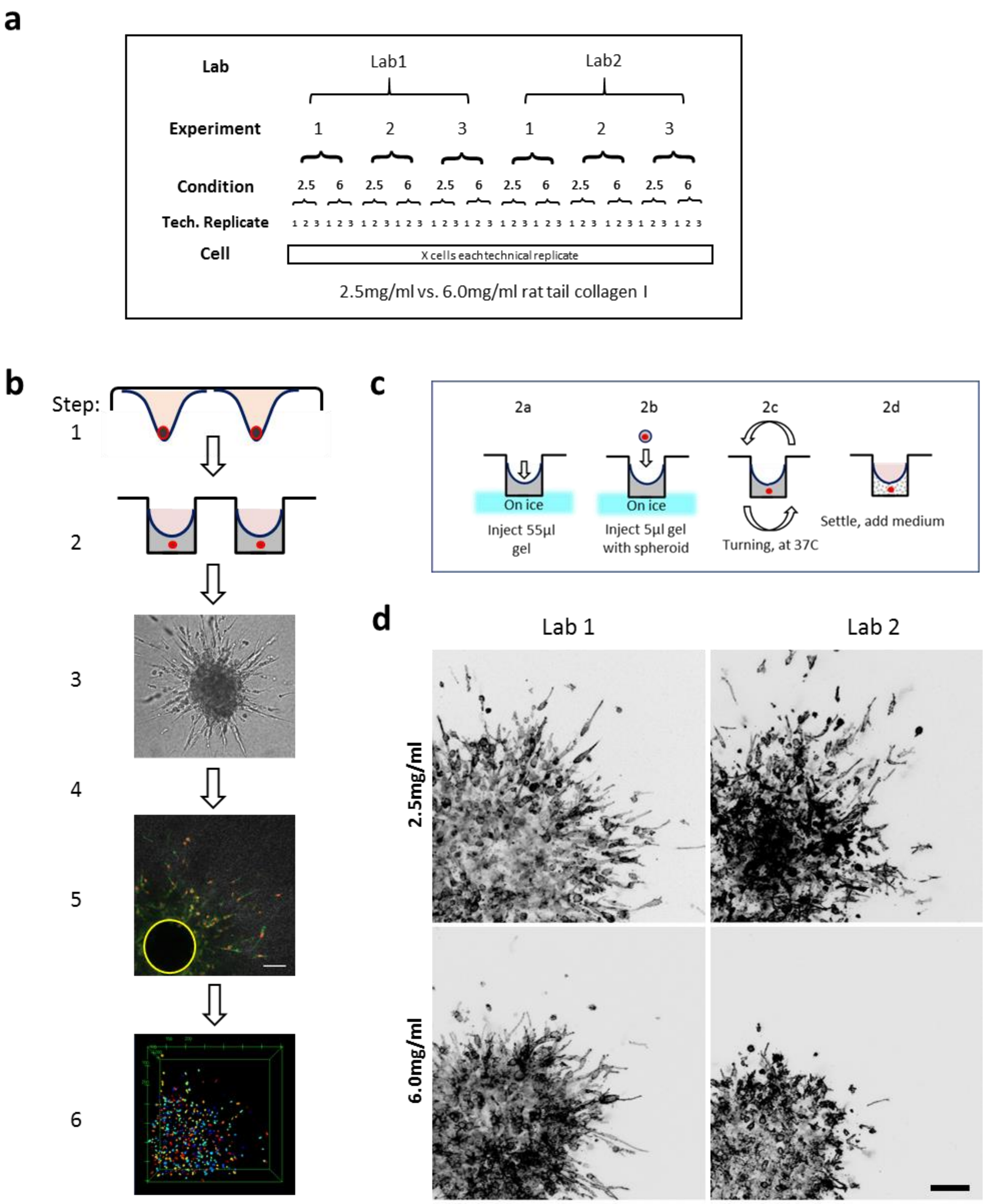


**Supplementary Figure 7. Study design and initial results of the 3D cell invasion experiment. a. Schematic of the study design,** resulting in collagen density dependent 3D invasion**.** The study involved two independent laboratories, three independent experiments in each laboratory, two conditions (2.5 mg/mL or 6 mg/mL collagen) in each experiment, with three technical replicates in each condition. **b. Overview of the pipeline**. Step 1, creation of spheroids (hanging drop assay), 1000 cells/drop, HT1080 cells. Step 2, embedding in rat tail collagen, concentration 2.5 or 6.0 mg/ml, in 96-well imaging plate, 1 spheroid/60µl gel. Step 3, incubation for 24h, 37°C, 10% $CO_2$. Step 4, 4% PFA fixation and immunofluorescent staining. Step 5, confocal imaging, objective 20x/0.8NA, stack size 708x708x120um, voxel size1.2x1.2x2µm, channels: nuclear DNA (red), F-actin (green) and reflection (grey). Bar: 100µm. Example data represent HT1080 cells, N=3, 2.5 mg/ml. Step 6, 3D nuclear segmentation and quantification of nuclear migration distances from manually annotated spheroid core (yellow circle in step 5). Segmentation precision = 88.05% and recall = 91.00% (see materials and methods). **c. Individual procedures performed in step 2 of the assay (described in b).)**. **d. Sample images of the produced datasets by different labs.** HT1080 cells were seeded in the collagen condition 2.5 versus 6.0mg/ml. Bar: 100µm**.**

**Supplementary Fig. 8**

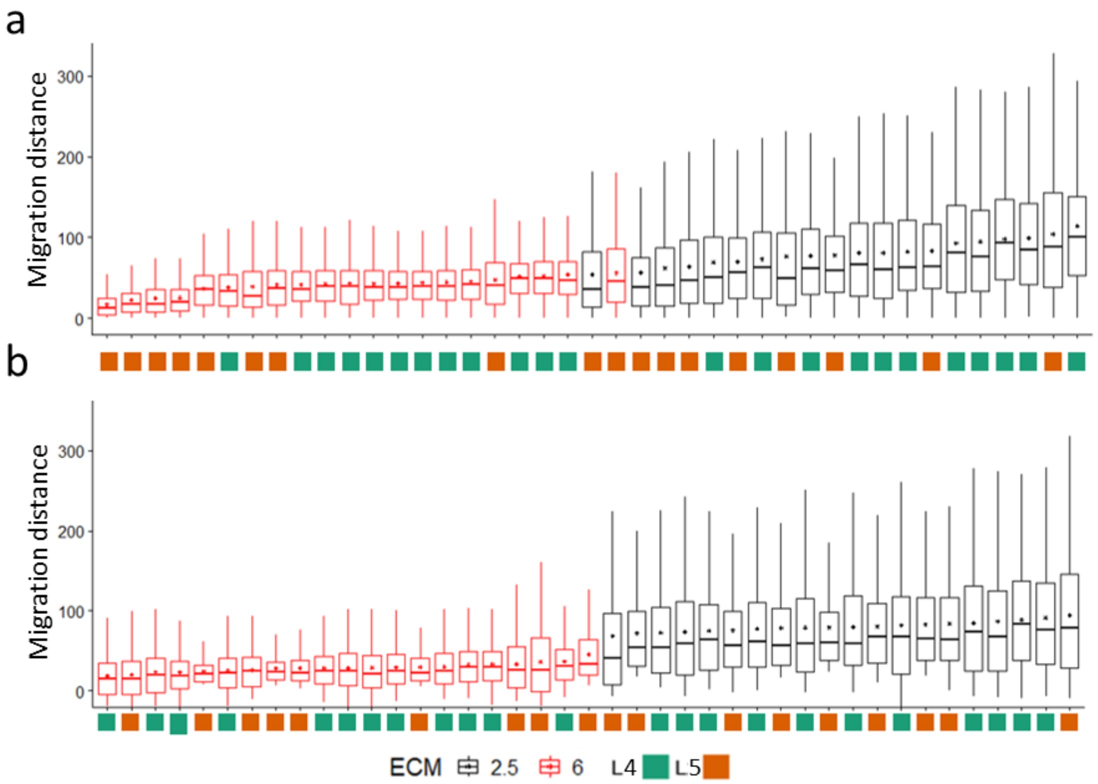


**Supplementary Figure 8**. **3D cell migration distance before (a) and after (b) batch effect removal.** Results are sorted in each case by increasing median order. Each boxplot shows the migration distance of a technical replicate. Laboratory #4 and laboratory #5 are shown with green and orange color respectively below the boxplots. ECM 2.5 mg/mL and 6 mg/mL are shown with black and red color, respectively. For the boxplots, in each box, the central mark indicates the median, and the bottom and top edges of the box indicate the 1st quartile and 3rd quartile respectively. Lines above and below the boxes show the high and low ranges of the migration distance. The dot in each box indicates the mean value.

### Supplementary table 1. Cumulative variability definitions

| Description | Formula | Number of possible combinations | Dataset contains all observations that belong to |
|---|---|---|---|
| 2 Replicates | $L_i(P_i(E_i(R_{w_2})))$ | 3*3*3*3 = 81 | $L_1P_1E_1R_{1\&2}$<br>(…)<br>$L_3P_3E_1R_{2\&3}$ |
| 3 Replicates | $L_i(P_i(E_i(R_{w_3})))$ | 3*3*3*1 = 27 | $L_1P_1E_1R_{1\&2\&3}$<br>(…)<br>$L_3P_3E_3R_{1\&2\&3}$ |
| 2 Experiments | $L_i(P_i(E_{w_2}(R_{w_3})))$ | 3*3*3*1 = 27 | $L_1P_1E_{1\&2}R_{1\&2\&3}$<br>(…)<br>$L_1P_1E_{2\&3}R_{1\&2\&3}$ |
| 3 Experiments | $L_i(P_i(E_{w_3}(R_{w_3})))$ | 3*3*1*1 = 9 | $L_1P_1E_{1\&2\&3}R_{1\&2\&3}$<br>(…)<br>$L_3P_3E_{1\&2\&3}R_{1\&2\&3}$ |
| 2 Persons | $L_i(P_{w_2}(E_{w_3}(R_{w_3})))$ | 3*3*1*1 = 9 | $L_1P_{1\&2}E_{1\&2\&3}R_{1\&2\&3}$<br>(…)<br>$L_3P_{2\&3}E_{1\&2\&3}R_{1\&2\&3}$ |
| 3 Persons | $L_i(P_{w_3}(E_{w_3}(R_{w_3})))$ | 3*1*1*1 = 3 | $L_1P_{1\&2\&3}E_{1\&2\&3}R_{1\&2\&3}$<br>$L_2P_{1\&2\&3}E_{1\&2\&3}R_{1\&2\&3}$<br>$L_3P_{1\&2\&3}E_{1\&2\&3}R_{1\&2\&3}$ |
| 2 Labs | $L_{w_2}(P_{w_3}(E_{w_3}(R_{w_3})))$ | 3*1*1*1 = 3 | $L_{1\&2}P_{1\&2\&3}E_{1\&2\&3}R_{1\&2\&3}$<br>$L_{1\&3}P_{1\&2\&3}E_{1\&2\&3}R_{1\&2\&3}$<br>$L_{2\&3}P_{1\&2\&3}E_{1\&2\&3}R_{1\&2\&3}$ |
| 3 Labs | $L_{w_3}(P_{w_3}(E_{w_3}(R_{w_3})))$ | 1*1*1*1 = 1 | $L_{1\&2\&3}P_{1\&2\&3}E_{1\&2\&3}R_{1\&2\&3}$ |

Where
$i = 1,2,3$
$\underline{\mathbf{w_n}} = (w_1, w_2, \dots, w_n)$, an n-tuple of natural numbers such that
$\underline{\mathbf{w_n}} \in \binom{\mathbb{N}_3}{n}$ , the set of all combination of the elements of $\mathbb{N}_3$ taken in groups of $n$ where $\mathbb{N}_3 \equiv \{1,2,3\}$, the set of the first 3 natural numbers.
For instance, $n = 2 \Rightarrow \binom{\mathbb{N}_3}{2} = \{1\&2, 1\&3, 2\&3\}$ and $n = 3 \Rightarrow \binom{\mathbb{N}_3}{3} = \{1\&2\&3\}$
