## Supplementary material for "Multi-site assessment of reproducibility in high-content live cell imaging data": Protocol for HT1080 2-D Migration_Live Cell Imaging

**By Jianjiang Hu**

**Materials:**

**Cell line:** HT1080 labeled with Lifeact-mCherry & H2B-EGFP from Radboud (verified by CRICK). *use the cells for experiment already being passaged at least once after thawing and also before passage 20.

| Name | Company | Catalogue Number |
|---|---|---|
| high glucose DMEM | gibco | 41965-039 |
| FBS | gibco | 10270-106 |
| sodium pyruvate | gibco | 11360070 |
| penicillin/streptomycin | gibco | 15140-122 |
| DMSO | Sigma | D2640 |
| trypsin (10x) | Life technology | 15400-054 |
| T-75 flask | SARSTEDT | 83.3911.002 |
| T-25 flask | SARSTEDT | 83.3910.002 |
| 18G syringe needle | KDM | 900444 |
| 6-well plate | FALCON | 353046 |
| 96- Imaging Plate CG (Cover Glass) | Mo Bi Tec | 5241-20 |
| Collagen I | Corning | 354249 |
| Heat denatured 0.5% BSA | Sigma | A2153 |
| Y27632 (ROCK inhibitor) | BD | 562822 |

***Green labelled materials are provided by Staffan lab.**

**Cell culture medium**: 500 ml high glucose DMEM + 50 ml FBS + 5 ml sodium pyruvate + 5 ml penicillin/streptomycin (10000 U/ml).

**FBS free culture medium**: 500 ml high glucose DMEM + 5 ml sodium pyruvate + 5 ml penicillin/streptomycin (10000 U/ml).

**Trypsin (1×):** 5 ml trypsin (10×) + 45 ml PBS

**1mM Y27632**: Add 2.956 ml dd$H_2O$ into 1 mg Y27632 to get 1 mM solution, aliquot into 100 ul/tube and store in -20℃ (already done).

**Heat denatured 0.5 % BSA**: Make a 0.5% BSA/PBS solution (add 0.5 g of BSA to 100 ml of 1×PBS) and adjust pH to 7.40. Then heat solution in boiling water for 7-8 min (until it turns opaque). *If solution is over-heated it will turn into a solid gel! Therefore, after 7-8 min of heating, transfer the solution into ice and bring its temperature down to room temperature (already done).

**Preparation work:**

1. Thaw one tube of the HT1080-Lifeact-mCherry-H2B-EGFP cells in 37℃ water bath.
2. Transfer the cells to a 15 ml tube containing 9 ml of cell culture medium. Add slowly and dropwise.

3. Centrifuge at 240×G for 2 min.
4. Remove the supernatant and re-suspend the cell pellet with 5ml culture medium.
5. Transfer the cells and medium to a T-75 flask and add culture medium to 15 ml.
6. Culture the cells in 37℃ 5% $CO_2$ incubator.
3. When the cells reach 80-90% confluency, split the cells into three T-75 flasks.
4. Freeze the cells in 3 flasks into 30 cryopreservation tubes (1 ml/tube) for further use.

**Freeze cells in T-75 flask:**
1. Discard the culture medium and wash the cultured cells with 15 ml PBS once.
2. Add 2 ml trypsin into the flask and put in 37 ℃ incubator for 2-3 min until the cells detached from the surface.
4. Add 15 ml culture medium into the well and gently dissociate the cells with 1ml pipette.
5. Transfer the medium containing cells into a 50 ml centrifuge tube and centrifuge at 240×G for 2 min.
6. Remove the supernatant and re-suspend the cells with 5 ml culture medium.
7. Add 5ml freezing medium (20% DMSO in FBS) slowly into above cell suspension and mix uniformly.
8. Add above cell suspension into 10 cryopreservation tubes (1 ml/tube).
9. Put the tubes into the Mr. Frosty™ Freezing Container (or similar ones) and then in -80℃ fridge.
10. After 24 hs, transfer the tubes into -140℃ fridge for long term storage.

**Thaw the cells from stock:**
1. Thaw the cells in 37℃ water bath.
2. Transfer the cells to a 15 ml tube containing 9 ml of cell culture medium. Add slowly and dropwise.
3. Centrifuge at 240×G for 2 min.
4. Remove the supernatant and re-suspend the cell pellet with 5ml culture medium.
5. Transfer the cells and medium to a T-25 flask.
6. Culture the cells in 37℃ 5% $CO_2$ incubator.

**Routine cell culture:**

**Cell passage:**
1. Discard the culture medium and wash the cultured cells with 5ml PBS once.
2. Add 1 ml trypsin into the flask and put in 37 ℃ incubator for 2-3 min until the cells detached from the surface.
4. Add 5 ml culture medium into the flask and gently dissociate the cells with 1ml pipette.

5. Transfer the medium containing cells into a 15 ml centrifuge tube and centrifuge at 240×G for 2 min.
6. Remove the supernatant and re-suspend the cells with 1 ml culture medium.
7. Seed suitable amount (e.g. 1:10) of the cells into a new T25 flask containing 5 ml culture medium.
8. Culture the cells in 37℃ 5% $CO_2$ incubator.

Grow the cells in a T25 flask and passage when the cells reach ~80-90 % confluency. Do not let the cells grow to 100% confluency.

*Avoid everyday passage. e.g. If design the live cell imaging for 3 continuous days, split the cells in advance into 3 flasks with different initial cell number, and then use one of the flasks each day for the cell seeding in step 1.

**Live cell imaging:**

**1. One day before the experiment:**
1.0 Cells in T25 flask should reach ~70-80% confluency before experiment.
1.1 Discard the culture medium in T25 flask.
1.2 Add 1 ml trypsin into the flask and put in 37 ℃ incubator for 2-3 min until the cells detach from the surface.
1.3 Add 5 ml culture medium into the flask and gently dissociate the cells with 1ml pipette.
1.4 Transfer the medium containing cells into a 15 ml centrifuge tube and centrifuge at 240×G for 2 min.
1.5 Remove the supernatant and resuspend the cells with 1 ml **culture medium**.
1.6 Count the cell concentration with the Fuchs-Rosenthal Counting Chamber.
1.7 Seed $2x10^5$ HT1080+Lifeact-mCherry+H2B-EGFP cells in one well of the 6 well plate and add 2 ml culture medium.
1.8 Culture the cells in 6-well plate in 37℃ $CO_2$ incubator overnight and then use as described in 2.2.

**2. On the day for live cell imaging:**

**2.1 Plate coating:**
**2.1.0 Plate drying:**
2.1.0.1 If a new imaging plate is used for the experiment, skip this step.

2.1.0.2 When re-using an imaging plate already used before, take the plate out from the incubator and put it in the hood in the cell culture room for 1 h to ensure that the surface is dry.

**2.1.1 Collagen I dilution:**
2.1.1.1 Add 10 ul 8.94 mg/ml Collagen I into 884 ul PBS and mix well to obtain homogenous 100 ug/ml Collagen I.
2.1.1.2 Add 200 ul of the 100 ug/ml Collagen I into 800 ul PBS to obtain 20 ug/ml Collagen I.

**2.1.2 Plate coating:**
2.1.2.1 Add 100 ul of the 20 ug/ml collagen I solution into each of 6 wells of the " Imaging Plate CG (Cover Glass) (96 well plate)" with normal yellow tips.
2.1.2.2 Put the plate in the cell incubator at 37℃ for 2 h.

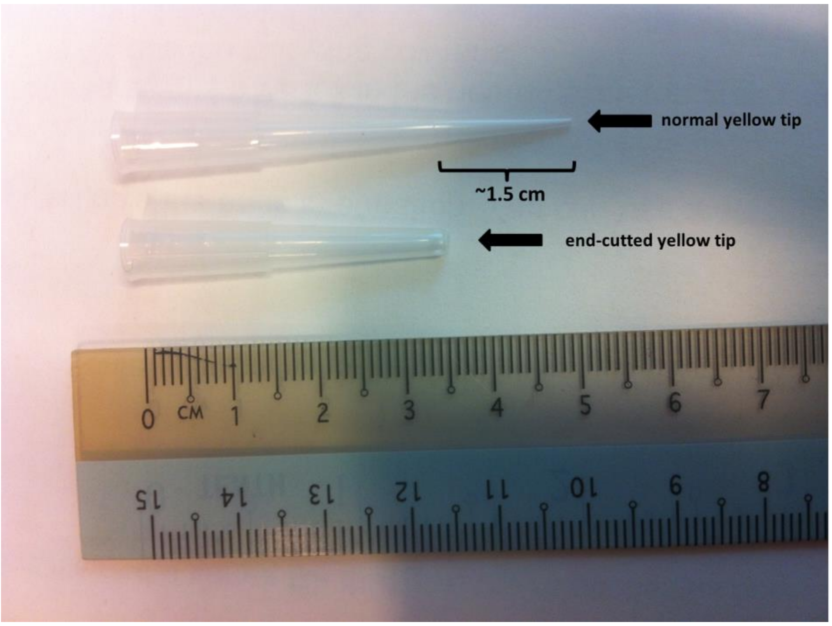


**Fig.1 Picture of the normal and end-cutted yellow tip**
Cut the sterilized yellow tip with sterilized scissors.
Sterilization of scissors: spray disinfectant (e.g. 70% ethanol or other disinfectant used in cell culture room) on to the scissors and then wipe with clean tissue paper.
*Alternatives: 1) Autoclave the scissors; 2) cut the yellow tips and then autoclave then in advance.

**2.1.3 Blocking:**
2.1.3.1 Flip the plate upside down and absorb the coating medium with tissue paper gently.
2.1.3.2 Then add 100 ul head denatured 0.5% BSA gently in each well with **end-cutted yellow tips**.

** **Burst the big bubbles remaining in the well with a heated 18G syringe needle carefully, and do not touch the bottom of the wells**
2.1.3.4 Put the plate at 37℃ minimum ~20 min (During which cell preparation (2.2) could be done).

**2.1.4 Washing:**
2.1.4.1 Flip the plate upside down and absorb the blocking medium with tissue paper gently.
2.1.4.2. Fill each well with 100 ul **FBS free culture medium** with **end-cutted yellow tips**.
2.1.4.3. Flip the plate upside down and absorb the washing medium with tissue paper gently.
2.1.4.4 Fill each well with 100 ul **FBS free culture medium** with **end-cutted yellow tips**.
****After each of 2.1.4 steps, burst the big bubbles remaining in the well with a heated 18G syringe needle carefully, and do not touch the bottom of the wells.**

**2.2 Cell preparation:**
2.2.1 Discard the culture medium in 6-well plate prepared one day before and wash the cultured cells with 2 ml PBS once.
2.2.2 Add 0.2 ml trypsin into the well and put in 37 ℃ incubator for 2-3 min until the cells detach from the surface.
2.2.3 Add 3 ml culture medium into the well and gently dissociate the cells with 1ml pipette.
2.2.4 Transfer the medium containing cells into a 15 ml centrifuge tube and centrifuge at 240×G for 2 min.
2.2.5 Remove the supernatant and resuspend the cells with 5 ml **FBS free culture medium**.
2.2.6 Centrifuge at 240x G for 2 min.
2.2.7 Remove the supernatant and resuspend the cells in 1 ml **FBS free culture medium**.
2.2.8 Count the cell concentration with the Fuchs-Rosenthal Counting Chamber.
2.2.9 Prepare 1ml cells at the concentration of $5 \times 10^3$ cells/ml in **FBS free culture medium**.

**2.3 Cell seeding:**
2.3.1 Seed 100 ul cells prepared from 2.2 in each collagen coated well from 2.1.4 (which already contains 100ul FBS free culture medium in it) with **end-cutted yellow tips**.
2.3.2 Tap the culture plate in two vertical directions to make the cell seeding even (For

right handed person, the tap direction should towards the up and right while left handed person to up and left). * Making cell seeding even is important for the following selection of imaging acquisition position.

2.3.3 Put the plate in the cell culture hood for around 10 min to let cells sink and attach to the plate. DO NOT put the plate back into the 37 ℃ incubator immediately after the cell seeding in 2.3.2. This 10 min allows the cells to sink and attach to the plate surface without being disturbed by the medium turbulence caused by the temperature fluctuation.

2.3.4 Put the plate back into 37 ℃ 5% $CO_2$ incubator and incubate for 2.5 h.

**2.4 Drug preparation:**

2.4.1 Add 15 ul 1 mM Y27632 into 485 ul **FBS free culture medium** in an 1.5 ml Eppendorf tube.

2.4.2 Put the above medium, together with 500 ul **FBS free culture medium** (in another 1.5 ml Eppendorf tube), into the microscope incubation chamber half an h before imaging acquisition for pre-warming.

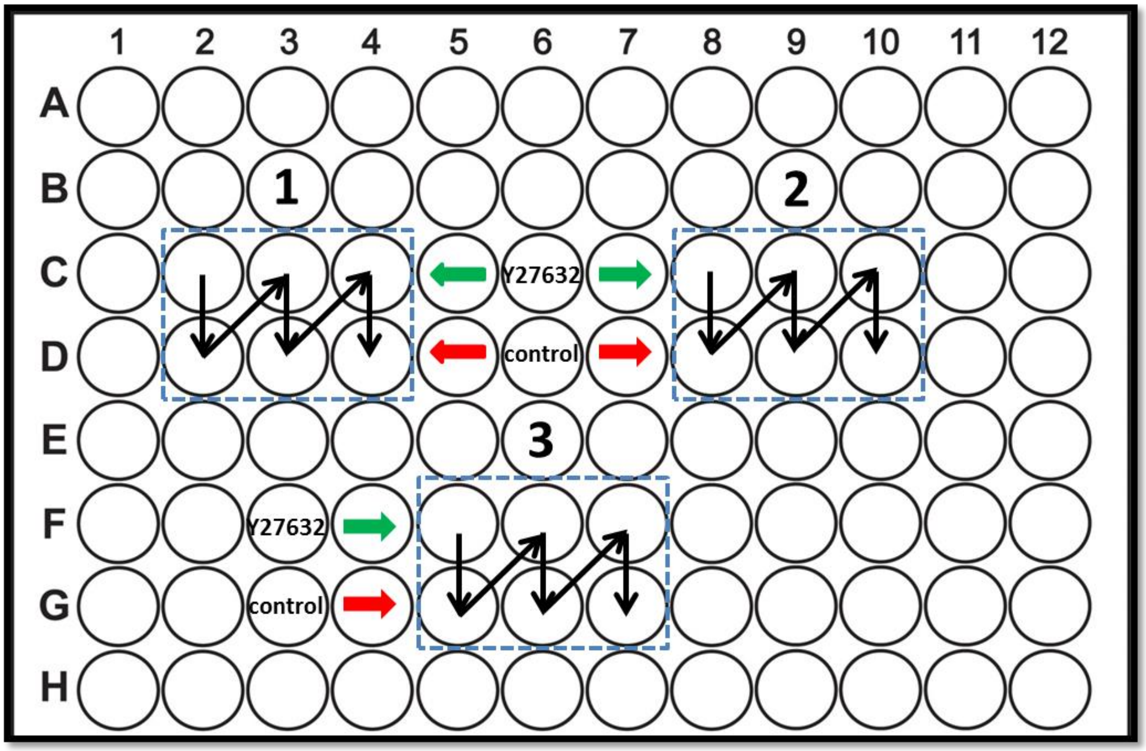


**Fig.2 Positions in the plate used for the experiment.**

**Blue rectangles** and the number above them show the 6 wells used for each of the three experiments. **Black arrows** show the acquisition sequence of live cell imaging. **Green**

**arrows** show three rows for Y27632 groups, while **red arrows** show three rows for control groups.

**2.5 Live Cell Imaging Setting**

2.5.1 Put the cell seeded plate from 2.3 on the microscope stage for environment stabilization, during which the microscope setting and position selection could be done (~0.5 h).

2.5.2 Microscope setting:

**Principles:** 1) With acceptable image quality for CellProfiler auto processing, excite the cells with as low laser power as possible to reduce the phototoxicity. 2) Open the pinhole to maximum. 3) Try to get one large image series in each well. 4) Time interval: 5 min; Total acquisition time: 6 h. 5) Use autofocus system to prevent focus drift during imaging. 6) Find suitable observation position (cell density similar with Fig. 3A) in each well (10× objective might help). Normally start from the center of each well. If too few or too much of cells in center, then look for nearby positions. But do not choose the position too close to the edge of well. 7) During image acquisition, avoid any light from the environment as much as possible (e.g. turn off the computer screen). 8) Also acquire bright field images if possible.

**Following are the microscope settings applied in Staffan lab:**

Confocal (Orion, Nikon A1) with 488 nm and 561 nm lasers, 20× objective (Plan Apo, NA 0.75), pinhole size: maximum (255.4 μm), 512×512 pixels (0.83 μm/pixel), 5×5 large image (2560×2560 pixels; 2125×2125 μm), zoom 1.5, no averaging

Nikon A1plus

{Scanner Selection}: Galvano
{Detector Selection}: DU4
{Optical Path Mode}: Manual
{First Dichroic Mirror}: 405/488/561/640
{First Filter Cube}:
{Second Filter Cube}: 525/50
{Third Filter Cube}: 595/50

| | | |
|---|---|---|
| EGFP Channel | {Laser Wavelength}: 488.0 | {Laser Power}: 0.3% |
| | {PMT HV}: 90 | {PMT Offset}: -2 |
| mCherry Channel | {Laser Wavelength}: 561.6 | {Laser Power}: 1.0% |
| | {PMT HV}: 100 | {PMT Offset}: -2 |
| TD | {PMT HV}: 100 | {PMT Offset}: 0 |

{Pinhole Size (um)}: 255.43
{HV LinearCorrection}: Off
{Scan Direction}: One way
{Scanner Zoom}: 1.499
{Scan Speed}: 1
{Channel Series Mode}: None
{Line Skip}: None
{Frame Skip}: 0
{Line Average Mode}: None
{Line Average/Integrate Count}: 0
Stim1 {Type}: Nothing
{Scan Speed}: 1
Stim2 {Type}: Nothing
{Scan Speed}: 1
Stim3 {Type}: Nothing
{Scan Speed}: 1

(**561 nm laser is specially adjusted on our system, should be ~0.2% on conventional confocal microscope**)
If possible, compare the power of 488 nm and 561 nm lasers with the measurements provided in the attached table with 10x objective and then adjust the laser powers used in the experiment accordingly. **It is necessary to use as low laser power as possible for the live cell imaging. Because the high laser power would be harmful for the cells.**

**Large image**: find the center of each well and take 5×5 images based on the centers, stitch with no overlap to get the final large image (2560×2560 pixels). If the acquisition software does not have the function of stitch large image, please provide the information of the positions of each small images for further stitching work with other software.
**Autofocus**: on

**Timelapse**: time interval: 5 min; total time: 6 h

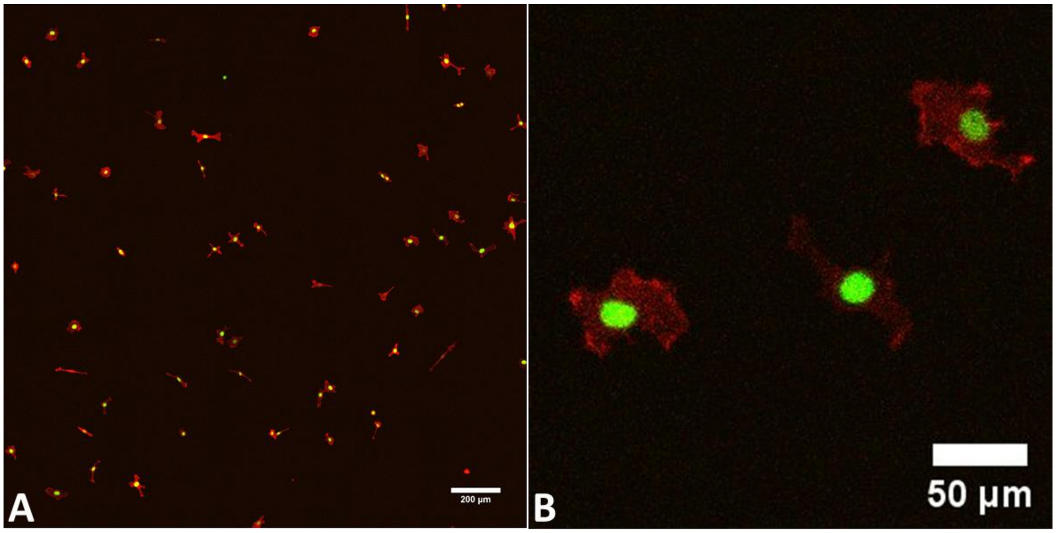


**Fig.3 Sample Images**

A) Full Image (2125 um × 2125 um, scale bar 200 µm). B) Zoom in.

(Brightness and contrast were adjusted for better display)

Please also see the following KI box link for sample movies:

https://ki.box.com/s/04x31gtkb359t88sjljtw3ss9i6yv1ay

**2.6 Drug addition:**

2.6.1 Add the drug just before imaging starts.

2.6.2 Add 100 ul medium containing (3 wells)/not containing Y27632 (3 wells) into the wells separately with an end-cutted yellow tips.

**2.7 Start the acquisition**

**2.8 After acquisition**

2.8.1 Take the imaging plate out and put it in the cell incubator for further usage.

2.8.2 Save data and turn off the microscope.

**2.9 Organization of acquired data**

Each person should have one folder, in which there are three folders containing three experiment repeats.

As the data from the timelapse are large, they are not suitable for online transfer. So please copy all the data from same lab into one hard drive with clear annotation and send to Staffan lab by mail. Thanks!

**Mail address:**

Jianjiang Hu
Clinical Molecular Biology / Lab Staffan Strömblad
Karolinska Institutet
Department of Biosciences and Nutrition
Hälsovägen 7-9
SE-141 83 Huddinge
Sweden

**Laser power measurement on Orion with 10x objective, values in uW. Low output 561 10 mW**

| Date | Laser % | 0 | 5 | 10 | 15 | 20 | 25 | 30 | 35 | 40 | 45 | 50 | 55 | 60 | 65 | 70 | 75 | 80 | 85 | 90 | 95 | 100 |
|---|---|---|---|---|---|---|---|---|---|---|---|---|---|---|---|---|---|---|---|---|---|---|
| 12-May-2016 | 488 nm | 2,628 | 88,67 | 176,6 | 268,1 | 356,0 | 444,3 | 532,9 | 621,0 | 708,8 | 796,9 | 883,2 | 975,0 | 1062, | 1149, | 1234, | 1319, | 1403, | 1493, | 1587, | 1664, | 1741,973 |
| 12-May-2016 | 561 nm | 1,338 | 125,3 | 251,2 | 372,9 | 500,4 | 626,4 | 745,7 | 874,7 | 999,3 | 1123, | 1248, | 1369, | 1498, | 1623, | 1749, | 1868, | 1990, | 2112, | 2240, | 2370, | 2490,843 |
