## Supplementary material for "Multi-site assessment of reproducibility in high-content live cell imaging data": 3D spheroid culture protocol

### Appendix I. Protocols for standardized 3D spheroid culture

*Preparation of Methylcellulose 12 mg/ml (5x):*
Weigh 6 g of methyl cellulose (Sigma, Cat.: M6385) in a 500 ml glass bottle, add a magnetic stir bar and autoclave the bottle. Add 250 ml of DMEM (Dulbecco Modified Eagle Medium, High Glucose, Gibco/Thermo Fischer Scientific, 10938-025) to the autoclaved methyl cellulose and incubate for 15 min at 60°C (shake the bottle to destroy clumps of methyl cellulose). Stir on a magnetic stirrer for 20 min, then add another 240 ml of DMEM, and continue stirring for 4 hours at room temperature. Keep the bottle overnight at 4°C. Add 10 ml of Penicillin-Streptomycin stock solution (10,000 U/mL, Gibco/Thermo Fischer Scientific, 15140122) and stir thoroughly. Centrifuge for 20 min. at 4000 rpm, at 4°C and carry the supernatant into 50 ml Falcon tubes.

*Cell culture:*
Human wild-type HT1080 fibrosarcoma (ACC315; DSMZ Braunschweig) are cultured (37°C at 10% $CO_2$ humidified atmosphere) in medium (DMEM, High glucose, Gibco) supplemented with 10% of fetal calf serum (FCS; Sigma Aldrich, F7524), Penicillin-Streptomycin (100 U/mL, Gibco/Thermo Fischer Scientific, 15140122), L-glutamine (2 mM) and sodium pyruvate (1mM; both Invitrogen) in a T75 culture flask. Grow cells up to 85-90% confluency before using.

*Generation of 3D spheroids (hanging-drop culture [1]):*

*This protocol describes the generation of multicellular spheroids of HT-1080 cells. These cells require supplemental collagen for proper aggregation.*

- Check cells in T75 flask for correct confluency (85-90%);
- Remove medium (as prepared for cell culture) and wash adherent cells with 1x PBS (room temperature);
- Detach cells with 2.5 ml of 2 mM EDTA (Invitrogen/Thermo Fischer Scientific, 15575020) in PBS for 10 minutes in a 37°C incubator;
- Check if the cells detached, mix the cell suspension with 7,5 ml medium and transfer it to a 15 ml tube;
- Spin down for 5 min at 1000 rpm;
- Remove the supernatant;
- Resuspend the pellet in 1 ml of medium and count the cells;
- Create a $2*10^6$ cells/ml suspension by addition of medium;
- Add 100 µl of cell suspension to a 15 ml tube;
- Add 1 ml methylcellulose 5x (end concentration 4 mg/ml);
- Add 2.6 µl of rattail collagen type I from stock (9.59 mg/ml, Corning, 354249, lot 6102001) to obtain a final concentration of 5 µg/ml of collagen in the suspension;
- Add 3.9 ml medium to have 5 ml of cell suspension with 40.000 cells/ml;
- Create 25 µl drops with a 200 µl multichannel pipet in the (hydrophobic) lid of a 15 cm culture dish (Greiner Bio-one, 639160);
- Put the plastic dish upside-down on the lid with the drops and gently turn the dish.
- Add 5 ml PBS 1x to the dish to avoid dehydration of the droplets;

- Incubate overnight at 37°C with 10% $CO_2$;

<u>*Spheroid embedding in 3D collagen:*</u>

*This protocol describes the embedding of HT1080 multicellular spheroids in rattail collagen I, in 96 well plates, using 1000 cells per spheroid, 1 spheroid per gel, in rattail collagen of 2.5 or 6 mg/ml. The protocol has been based on the protocol of Wolf et al., 2013 and Haeger et al. 2014, which describe spheroid embedding in low number of drop gels for different collagen concentrations [2,3].*

*The protocol aims at embedding approximately 18 spheroids per rattail collagen concentration. The successfully embedded spheroids will be divided into 3 groups with at least 3 technical replicates per group, containing a medium control, a vehicle control such as DMSO and a molecular interference such as a FAK inhibitor.*

*The embedding process consists of two steps: first, gel droplets without spheroids are deposed into the middle of the wells of a 96-well plate. Second, before polymerization of the gel, a small droplet of gel containing a spheroid is precisely pipetted into the middle of the gel in each well. The advantage of this protocol is that the position of the spheroids can be better controlled compared to direct deposition of a complete gel with a spheroid. Furthermore, all spheroids will sink down to the bottom of the gel droplet and therefore their position with respect to each other is synchronized in time. As a result, spheroid height with respect to the bottom of the plate can be tuned by applying a proper turning sequence during the polymerization process.*

*Use only one set of pipettes which have been properly calibrated! Deviations in the volumes can strongly influence the consistency and polymerization time of the gel, resulting in altered spheroid invasion phenotypes and bad control over spheroid positioning. Start this protocol with the 2.5 mg/ml concentration and continue with 6 mg/ml in the same plate, after polymerization of the first concentration.*

- Aspirate PBS form the hanging drops dish;
- Check the spheroids in the hanging drops by putting the dish with lid side down on a microscope. Mark the good (round, compact) spheroids;
- Cool the 96-well imaging plate *(µ-Plate 96 Well, cell culture surface coating, sterile, Thermo scientific NUNC #165305)* and 5x 1.5 ml pointy Eppendorf tubes on ice;
- Harvest the spheroids: take out the wrong spheroids with P1000 pipet with blue tip and rinse 32 good spheroids from the inclined lid with 5 ml PBS. Transfer the PBS with the spheroids to a 15 ml tube;
- Let the spheroids sink to the bottom of tube (about 3-5 minutes). Aspirate carefully and take care that there is 20 µl left at the pellet of the spheroids;
- Add 3 ml PBS and gently invert the tube 5x times (a wash step);
- Let spheroids sink to the bottom again and aspirate carefully;
- Add 3 ml medium and let the spheroids collect at the base of the tube;
- Start preparing the gels:

*Prior to addition of the collagen solution, a reagent mix of 10x PBS, 1N NaOH and MilliQ is prepared and divided in a part to be used for preparation of the first gel (step 1) and a part for the preparation of the second gel containing the spheroids (step 2). To improve control over the consistency of the gel, collagen solution is added to the reagent mix of each step just before deposition of the gel in the wells. The amount of NaOH added is carefully determined beforehand so that the final prepared collagen gels have the correct pH. Monitoring pH and keeping to the temperature in the protocol below is essential to obtain collagen gels with reproducible density (pore size).*

- Prepare a reagent mix for one collagen gel concentration in one cooled Eppendorf tube; Use the calculated reagent mix volumes for current RT collagen stock:
  *Current collagen stock: Corning Collagen Type I, ref 354249, lot 6102001, 9.59 mg/ml*

| | 2.5 mg/ml mix 5x 385 µl (for 6 gels) | 6.0 mg/ml mix 5x 385 µl (for 6 gels) |
|---|---|---|
| PBS 10x | 154 µl | 154 µl |
| NaOH 1N | 10 µl | 24.1 µl |
| Milli Q | 874.15 µl | 157.55 µl |

**Start with 2.5 mg/ml and then use calculations for 6mg/ml later on in the protocol**

- Distribute 1/5 of the mix over each of the 4 remaining Eppendorf tubes on ice (207,63 µl for 2.5 mg/ml and 67.13 µl for 6.0 mg/ml). Discard the tube with the remaining mix;
- Prepare the first gel without the spheroids: take up collagen very slowly (100.36 µl per ep for 2.5 mg/ml and 240.88 µl for 6.0 mg/ml) and add it to 3 of the 4 Eppendorf tubes on ice. Mix it thoroughly (10x up and down). *Use a P1000 pipette;*
- Add 77 µl of medium <u>without spheroids</u> to the same 3 Eppendorf tubes with mix and collagen solution. *The liquid collagen gel should appear transparent to very light yellow. Use a P200 which has been cut to increase the opening diameter;*
- Add the 3 mixes together in one of the 3 Eppendorf tubes and throw the remaining 2 tubes away. *Keep the gel on ice!*
- Aliquot the first gel into 18 wells: while the plate is on ice, pipet 55 µl gel in the middle of the wells. Avoid air bubbles;
- Check the pH of the remaining gel with pH paper (Merck Millipore pH 5.5-9.0 Neutralit). 2.5 mg/ml should be between 7.0 and 7.5 and 6.0 mg/ml should be near 7.0. *Note: a wrong pH strongly influences polymerization time and the consistency of the gel and therefore the position of the spheroids in the gel after polymerization!*
- Prepare the second gel containing the spheroids: add rattail collagen to the last Eppendorf tube on ice (100.36 µl for 2.5 mg/ml and 240.88 µl for 6.0 mg/ml). *Use a P1000 pipette with a blue tip*;
- Add 77 µl medium <u>with spheroids</u> to tube and mix gently and thoroughly (pipet 10x up and down). *Use a P200 with a cut tip to increase the opening diameter;*
- Take out 5 µl gel containing a single spheroid and deposit the volume in the middle of the gel, close to the bottom of the well. The total gel volume will be 60 µl per well. *Keep the tube containing the spheroids on ice while adding spheroids to 96 well plate!*
- Complete the 18 wells for 2.5 mg/ml;

- Polymerize the gels in such a manner that the spheroids remain in three dimensions, not touching any surfaces, while remaining within the working distance of the imaging objective to be used:

  2.5mg/ml:

  - Wipe the base of the plate (to remove ice);
  - Start the timer. Place the plate for 5 min 15 s in the incubator the correct way up (to allow all spheroids to sink to the bottom). *Harvest the spheroids for the 6.0mg/ml condition;*
  - While in the incubator, invert the plate every 60 seconds until polymerized (The gel will appear milky, 14-15 minutes on the timer);
  - After polymerization, leave the plate in normal position in the incubator until 18 minutes have passed on the timer; *Prepare the reagent mix without collagen solution for the 6 mg/ml gel;*
  - Place the plate back on ice during further preparation of the 6mg/ml gel;

  6.0 mg/ml:

  - Start the timer. Keep plate 4 minutes on ice (all spheroids sink to bottom);
  - Wipe the base of the plate (to remove ice);
  - Place the plate upside-down in the incubator for 2.5 minutes;
  - While in the incubator, invert the plate every 60 seconds until polymerized (~9 minutes on the timer);
  - After polymerization, leave the plate in normal position in the incubator until 30 minutes have passed on the timer. *Prepare the media for the different conditions in the plate. Keep the media at 37*°C;

- After polymerization of both collagen concentrations, check embedded spheroids and divide the good spheroids into 3 groups, randomly distributed over the plate: for example, Medium, DMSO and a pharmacological inhibitor;
- Make sure every group contains at least 3 successful technical replicates (briefly inspect spheroids by microscope);
- Add 175 µl medium to each well: add medium with pharmacological inhibitor (e.g. FAK inhibitor, TOCRIS PF573228, 2 µM work concentration in medium), medium with a vehicle control (e.g. in case of FAK inhibitor, DMSO) and medium only to the wells;
- Take time = 0 snapshots of each spheroid with a brightfield microscope;
- Incubate 24 hours at 37°C to establish cancer cell invasion in three dimensions, prior to fixation.

*Calculation of reagent quantities for preparation of a rattail collagen gel:*

- *Variables, chosen by experimenter:* $V_{final}$ (final volume, µl), $C_{stock}$ (stock concentration of the collagen solution, mg/ml), $C_{final}$ (final collagen concentration of the gel, mg/ml) and *p* (calibration factor to establish the correct pH of the collagen gel).
- *Reagent quantities to be calculated:* mQ (milliQ water, µl), Cells (medium with cells, µl), NaOH (solution of potassium hydroxide, 1N, µl), Coll (Collagen stock solution, µl) and PBS10x (10x PBS solution, µl).
- *Formulas:*

  $Coll = C_{final} * V_{final} / C_{stock}$

Cells = 0.2 * $V_{final}$
NaOH = *p* * Coll
PBS10x = ($V_{final}$ - Cells) / 10
mQ = $V_{final}$ – Cells – PBS10x – Coll – NaOH

The factor *p* was determined upon arrival of a new batch of rattail collagen solution, which has its own unique stock concentration and material properties. The factor was derived empirically by adjustment of the pH of a prepared collagen gel to 7.4, using the previously described pH measurement paper. *Starting point: p = 0.02.*

*Fixation and immunofluorescent staining of spheroids in a 3D collagen matrix:*

- Inspect the embedded spheroids under microscope to judge if the spheroids display a 3D invasion phenotype. Spheroids that display significant migration along 2D surfaces should be discarded;
- Remove the medium from the wells with multichannel pipette;
- Add 200 µl PBS;
- Remove the PBS and repeat this step;
- Add 200 µl 4% PFA per well and incubate for 15 minutes at room temperature;
- Remove PFA solution and wash 4x 15 minutes with 200 µl PBS at room temperature;
- Permeabilize and block with 200 µl per well labeling solution (see below) for 1 hour, at room temperature;
- Dilute primary antibodies in labeling solution;
- Remove all liquid from each well, gently!
- Add 150 µl primary antibodies in labeling solution to each well and incubate overnight on a shaker at 4°C;
- Remove almost all liquid from each well, gently!
- Wash 5 x 15 minutes with 200 µl PBS + 0.1 % Tween;
- Dilute secondary antibodies, Phalloidin 633 and DAPI in labeling solution;
- Remove PBS + 0.1% Tween from wells and add 150 µl of secondary antibodies in labeling solution to each well;
- Incubate for 4 hours on a shaker at room temperature, in the dark;
- Wash 5x 15 min with PBS + 0.1% tween 200 µl;
- Remove PBS and add 200 µl PBS with 0.05% azide to each well;
- The plate must be stored (preferably <48 hours) at 4°C prior to imaging.

Antibodies and labeling reagents:

- DAPI (Sigma, D9542): stock concentration at 1 mg/ml in water. Stain with 2 µg/ml. Dilute in steps to prevent precipitation;
- AlexaFluor633-Phalloidin (Molecular Probes, A22284): apply at a 1:200 dilution;
- Primary antibody YAP, rabbit mAb IgG (D8H1X, Cell Signaling Technology, #14074): stock concentration 5.5 µg/ml. Apply at a 1:200 dilution (dilute to 0.0275 µg/ml);
- Isotype control, rabbit mAb IgG (DA1E, Cell Signaling Technology, #3900): stock concentration 2.5 mg/ml. Dilute to 0.0275 µg/ml;

- Secondary antibody, highly cross absorbed goat-ant-rabbit IgG1 AlexaFluor488 (Thermo Fischer Scientific, A11034 2017, 1:1 in glycerol): stock concentration 2 mg/ml. Apply at a 1:200 dilution (dilute to 10 µg/ml);
- 10% BSA solution is freshly prepared from BSA powder (Fraction V, Biomol) in MilliQ.
- Labeling solution is freshly prepared from 0.3 % Triton X-100 in PBS, 10 % normal goat serum and 1 % BSA (from freshly prepared 10 % BSA solution);
- PFA 4 % solution is freshly prepared from frozen (-20°C) PFA 8 % stock solution (PFA dissolved in mQ, which was adjusted with 1N NaOH until the suspension became clear). PFA 8 % is thawed and the suspension is dissolved in a 60°C water bath. An equal volume of 0.2 M Phosphate buffer is added to reach a concentration of 4 % PFA.

*Adaptations to the standardized protocols:*
*To apply the standardized protocols to other cell lines and organoid culture, the following adaptations and additions were made to the hands-on protocols described above:*

Spheroid assay with MDA-MB-231 cell line:
- In order to generate 3D spheroids, a double concentration of methyl cellulose (4.8 mg/ml) and addition of 10 µg/ml Purecol Bovine Collagen Solution (Type I, 3 mg/ml, Cell Systems, cat. Nr. 5005-100ML) instead of rattail collagen was used;
- A Spheroids were incubated for 48h to establish a clear invasion phenotype.

Adaptation spheroid assay to CRC organoids (P19TA, #9 neon, #14):
- *Adaptation to protocol for cell culture:* human lung MRC-5 fibroblasts were cultured with the following modifications to the cell culture protocol: cells were maintained in an atmosphere with 5% $CO_2$. Cells were cultured up to full confluency before use, until a bundled structure of cells was observed.
- *Organoid culture:* the patient-derived CRC organoids were established as part of a living biobank of CRC10, and were kindly provided by the HUB foundation (hub4organoids.eu). Organoids were cultured in 70% Matrigel (Corning, 356231) with basal medium 2+ (BM2+) containing advanced-DMEM/F12 (Life Technologies, 12634028), supplemented with penicillin (100 U/ml) and streptomycin (100 µg/ml; PAA), HEPES (10 mM, Lonza, BE17-737E), GlutaMAX (400 µM, Life Technologies, 35050038), B27 (0.2X, Life Technologies, 17504044), N-Acetyl-L-cysteine (1 mM, Sigma-Aldrich, A9165-5G), Noggin (10%), A83-01 (500 nM, Biovision, 1725-1) and SB202190 (10 µM, Sigma-Aldrich, S7067) at 37°C and 5% $CO_2$.
- *Adaptations to protocol for generation of 3D spheroids:* organoids were directly harvested from culture, after an incubation period of approximately 12 days. At this timepoint the organoids approached a comparable size to the HT1080 spheroids (approximately 300 µm, for spheroids of 1000 cells).
- *Adaptations to protocol for Spheroid embedding:* in order to harvest the organoids, Matrigel was dissolved with ice cold PBS and organoids are transferred to a 15 ml falcon tube, followed by three washing steps with 10 ml ice cold PBS and centrifugation in between (5 min, 111 g, 4°C). Organoids were resuspended and kept on ice in 300 µl BM2+ medium until embedding in collagen.
  Then, organoids were embedded in a collagen matrix co-cultured with MRC-5 fibroblasts in the following manner: first, MRC5 cells from on-plastic culture (80% confluency) were

detached using Trypsin /EDTA (0.075% /2 mM, Thermo Fisher Scientific) in PBS followed by addition of medium containing 10% FCS. Second, cells were spun down, resuspended in BM2+ and counted. Then, during preparation of the collagen mixture for the first embedding step, the addition of 77 µl of medium was replaced by the addition of 40 µl of BM2+ medium and 37 µl of MRC5 cell suspension (520,000 cells/ml). For the preparation of the collagen mixture of the second embedding step, the BM2+ medium containing the resuspended harvested organoids was used. *Standardization of the organoid height with respect to the bottom of the imaging plate was not possible due to the broad distribution in organoid size. The spheroids from the P19TA, #9 neon and #14 cell lines were embedded in respectively 6, 5, and 5mg/ml collagen.* For long-term culture, BM2+ medium, supplemented with 2.5 % FCS to induce fibroblast elongation and maintain cell vitality, was added. Medium was refreshed at day 3 and every second consecutive day.

*Fixation and immunofluorescent staining:* At day 11 after embedding, samples were fixed following the procedures above, except for the fixation time of 30 minutes at room temperature. The #9 and #14 organoids were stained with DAPI and Phalloidin633 only.

*Ontology terms, candidate for, or part of MIACME:*

- Cell culture, dimension: 3D (among other terms downstream of cell culture; e.g. 2D, 2/3D;)
- Cell culture, 3D: Spheroid culture (downstream of 3D, among other terms such as interface assay, sandwich assay, single cell assay, organoid culture, …);
- Cell culture: Co-culture (downstream of cell culture. It must also be possible to note down a second cell line when culturing with different cell lines at the same time);
- Cell culture, 3D, Spheroid culture, method: hanging drop (downstream of spheroid culture);
- Cell culture, 3D, Matrix: rattail collagen I (downstream of 3D, among other terms such as Matrigel, hydrogel);
- Pharmacological inhibitor;
- Labeling type: immunofluorescent (downstream from labeling, among others such as fluorescent reagent, non-fluorescent reagent, immunostaining, immune-gold, ...)
- Labeling type, fluorescent reagent: DAPI (among others like Hoechst, phalloidinAlexa633)
- Labeling type, immunofluorescent: YAP mAb;

*References:*

[1] D. Del Duca, T. Werbowetski, R.F. Del Maestro, Spheroid preparation from hanging drops: characterization of a model of brain tumor invasion, J. Neuro-Oncol. 67(2004) 295–303.

[2] K. Wolf, M. Te Lindert, M. Krause, S. Alexander, J. Te Riet, A.L. Willis, et al., Physical limits of cell migration: control by ECM space and nuclear deformation and tuning by proteolysis and traction force, J. Cell Biol. 201 (2013) 1069–1084.

[3] A. Haeger, M. Krause, K. Wolf, & P. Friedl, Cell jamming: Collective invasion of mesenchymal tumor cells imposed by tissue confinement, Biochimica et Biophysica Acta - General Subjects, 1840(8), (2014) 2386–2395. https://doi.org/10.1016/j.bbagen.2014.03.020
