## Supplementary material for "Multi-site assessment of reproducibility in high-content live cell imaging data": 3D spheroid microscopy protocol

### Appendix II. Standardized microscopy

*The following workflow defines the standardized acquisition of microscopy datasets from 3D spheroid cultures. The parameter sets are defined for point scanning confocal microscopy and can be adapted to other microscopy systems, in a platform- and vendor-independent manner.*

#### *General settings:*

- Sample positioning in the scan field: preferably place the spheroid in the lower left corner, in such a manner that the core border is touching the image border. This can be seen best in the reflection channel.
- Set the Z-range up to 120 µm, starting from invading cells above the spheroid towards roughly the middle of the core (from z= 1/2 to z= 4/5 of spheroid dimensions).
- Detection range: 8 bit.
- Set the following for the optical path and detection configuration:
    - Sequential scanning, to record spectrally overlapping labels as well as reflection and transmission signal independently.
    - Scan 1: 488 nm (YAP) and 633 nm (F-actin; Phalloidin633).
    - Scan 2: 405 nm (DAPI) and 561 nm (reflection and transmission)
- The laser power and amplification (voltage) should be set in such a manner, that the brightest relevant cells make use of the full detection range. It is no problem if cells near the glass or cells in 2D are overexposed. But, deeper cells should be visible with sufficient signal to background, especially in the DAPI channel which is used for segmentation.
- Set laser power as high as possible to get good SNR while scanning fastest, without saturation of the dyes (2x intensity=2x emission).

#### *Microscopy system-dependent settings*

##### 1. *Zeiss LSM880*

- Use the following microscope settings:
    - Objective: Zeiss Plan-Apochromat 20x/0.8NA
    - Scan field: 708.5 $µm^2$
    - Pixels: 608x608, 1.2 µm/pixel
    - Pixel dwell time: 1.3 µs (fastest) with bi-directional scanning (adjust x and y manually)
    - Multi line averaging: 3
    - Optional: increase laser intensity linearly with increasing scan depth in 3D sample to maintain mean fluorescence intensity with the weakest staining as reference. Preferably increase laser power over multiline. At these coarse resolution setting samples do not bleach easily and high laser power is tolerated.
    - Z-depth: 2 µm
- Filter sets for excitation beam path for scan2: 80/20 dichroic mirror, such that reflection can be detected clearly.
- Use the GaAsP detector for the detection of the immunolabeling of weak signals.

2. *Olympus FV1000*

- Use the following microscope settings:
    - Objective: Olympus UPLSAPO 20x air/0.75NA
    - Scan field: 621 $\mu m^2$
    - Pixels: 600x600, 0.994 µm/pixel
    - Pixel dwell time: 2 µs
    - Multi line averaging on 1-3, depending on the intensity of the weakest staining. Preferably increase laser power over multiline. At these coarse resolution setting samples do not bleach easily and high laser power is tolerated.
    - Z-depth: 2 µm
- Set the Z-range up to 120 µm, starting from invading cells above towards roughly the middle of the core (from z= 1/2 to z= 4/5 of spheroid dimensions).
- Set the following for the optical path and detection configuration:
    - Sequential 'virtual mode' scanning, to record one immuno-labelling, reflection and transmission signals: Only the laser lines are switching on detector settings are changed, no filter settings are being changed between scans.
- Filter sets detection channels: DM 405/488/559/635, DM490, DM560, Mirror; 430-470, 505-540, 575-675.
- Detectors to be used are: HV 600 (channel 1, DAPI), 600 (Channel 2, 488), 650 (channel 3, phalloidin) or 560 (channel 3, reflection), and 230 (transmission), set the gain to 1.

<u>*Ontology terms, candidate for, or part of MIACME:*</u>

- Pixel dimension X (downstream of Imaging, Image properties, among pixel dimension Y, Z);
- Type: confocal microscopy (downstream of Imaging, Imaging devices, Microscopy);
- Device name: Zeiss LSM880 (downstream of Imaging, Imaging devices);
- Objective: Zeiss Plan-Apochromat 20x/0.8NA (downstream of Imaging, Imaging devices);
- Channels: DAPI, YAP, phalloidin, reflection, transmission (downstream of Imaging, Imaging devices, Microscopy);
