## Supplementary material for "Multi-site assessment of reproducibility in high-content live cell imaging data": 3D spheroid image analysis protocol

### Appendix III. Standardized image analysis

*This workflow is implemented in Fiji as the Nucleus Annotation 3D (NA) and the Cell Migration Analyser 3D (CMA) plugin sets and was distributed to 3 independent labs (RUMC, CRICK, UGENT) for standardized analysis of independent datasets from spheroid culture performed in each lab independently. The plugins were distributed among the WP6 partners and the final version plus a more in-depth manual will be deposited at GitHub.*

*The complete workflow consists of two distinct phases. Both are given as separate workflows below. The first workflow is used to determine the optimal input parameters which will be used during the second workflow in which the image data set is analyzed.*

#### Optimize input parameters workflow

*The optimization workflow is used to determine what the best set of input parameters is for the plugin set on the image data set. This workflow should be performed only once per collection of image data, as long as a similar experimental and imaging set up is chosen. Any changes to the setup that affect image parameters such as average cell size, the amount and type of background noise or image resolution, for example, need a re-optimization of the input parameters.*

*For all steps in which the software is to be used during the workflow it is indicated whether these are to be applied by the operator of the software (O) or automatically performed by the software itself (A).*

1. *Select an optimization set*

- From the available images, select a representative set that will be annotated by hand and serve as a 'golden truth' during the optimization.
- Try and include at least one image per condition.
- Partial images are fine to use, as long as the part is rectangular in form and it is well defined which part of the image is to be used. Crop the image to size.
- Note that these images should ideally not be used in the actual data gathering after optimization. Please note that only the nucleus and actin signal is used for this purpose, so any images that may have flaws in other channels are a good choice for this set.

2. *Create a 'golden truth' annotation set*

The following steps are needed to create a golden truth for each image in the optimization set:

- Start the Nucleus Annotation 3D plugin. (O)
- Select an image file to annotate through the dialog. (O)
- If the file has been imported via the BioFormats dialog, confirm this when prompted. (O)
- If the image has multiple channels, select the channel number of the nucleus signal. (O)
- Duplicate the chosen channel. (A)
- Adjust the brightness and similar settings to improve visibility if needed and confirm when ready. (O)

- Annotate each nucleus in the image. Follow the dialog explanation of the steps to perform. (O)
  Steps 1, 2, and 3a are obligatory for parameter tuning on just segmentation. If migration mode analysis is to be used during measurements, either step 3b, 3c, or 3d is needed for every cell as well.
- When finished with all the nuclei in the image click 'Done'. (O)
- Save the annotation next to the image in a file with the same name but the '_corrections.txt' extension. (O)

3. *Test parameter settings against annotation*

For each annotated image in the optimization set, perform the following steps:

- Open the image. (O)
- Start the Marker Image Creator 3D plugin. (O)
- Select 'Manual points' in the dialog. (O)
- Find and select the annotated 'corrections' file for this image in the file chooser dialog. (O)
- Select the directory in which the marker files will be saved. (O)
- Create the sub-directories 'Marker_Files' and 'Marker_images' in the chosen directory if they do not exist yet. (A)
- Save the marker file and marker image. (A)
- Use the Marker Image Creator 3D plugin as described in the Image Analysis Workflow. (O)
- Use the following as initial input parameters:
  - To find the minimal and maximal nucleus size parameters, measure some nucleus sizes by hand using the straight-line ROI tool:
    - Find the slice on which the nucleus is at its largest.
    - Draw a line ROI from one side of the nucleus to the other and note the distance displayed on the FIJI main dialog. Measure across an average part of the nucleus here, so try to avoid any dents or bulges.
    - Measure both nuclei that are smaller and larger to determine a good lower and upper boundary for the nucleus size. Once again, avoid outliers and instead focus on nucleus sizes that are common but on the low or high end of the scale.
    - 3 or 4 nuclei of a smaller size should suffice to find an average minimal nucleus size and a similar number of larger nuclei will be needed for the maximum nucleus size.
  - Subtract the higher size value from the lower value and halve the difference for the step size (round up to one decimal behind the decimal point). This will result in three steps during the LoG filtering.
  - For Noise, use the value of 1.
  - As Minimum Value, use ⅓ of the 'Maximum size of nucleus' parameter.
  - The RadiusXY value should similarly start at ⅓ of the 'Minimum size of nucleus'.
- The segmentation needs a choice of either a 3D mean or 3D median filter followed by a threshold. Determine a combination that looks good by eye as your initial parameters. One combination is needed for the nucleus channel and one for the actin channel. (O)

- Use the Marker Controlled Watershed 3D plugin as described in the Image Analysis Workflow below with the initial parameter combinations from the previous step. (O)
- Use the Feature Extraction 3D plugin as described in the Image Analysis Workflow below. (O)
- Review the results and retry the plugins with different parameters until satisfied. (O)

#### Image Analysis Workflow

The image analysis workflow describes the order in which the plugins are to be used per image. For each plugin, it gives the name of the plugin (with the plugin set in parentheses) and the individual workflow for that plugin. For all the plugin-specific workflow steps it is indicated whether these are to be applied by the operator of the software (O) or automatically performed by the plugin (A).

##### *1. Spheroid Annotation 3D (NA)*

- Load the image. (O)
- Identify the channel to annotate on. Usually the reflection channel. (O)
- Duplicate the chosen channel. (A)
- Adjust the channel brightness if desired. (O)
- Select a slice in which the spheroid outline can clearly be identified. Preferably a slice closer to the centre of the spheroid. (O)
- Click to add an annotation point on the edge of the spheroid. Select three points spread out along the edge. (O)
- Once three points have been selected, draw a circle through the three points. (A)
- Adjust the three points (by removing and adding again) until the circle best fits the spheroid edge. (O)
- Select another slice in which the spheroid edge is clearly identifiable. This slice should be considerably higher or lower than the previous slice and still contain an open spheroid edge (i.e. not the top or bottom slices of the spheroid). (O)
- Add one point to the edge of the spheroid. (O)
- Draw a sphere through all four annotation points. Adjust the z-size of the sphere to take the X/Y versus Z resolution ratio account. (A)
- Go through the image to judge the fit of the sphere to the spheroid edge. Adjust if necessary by removing and adding the annotation points. Note that the first slice should always have three points and the latter slice always just one. The one point is best for adjusting the sphere's size, while the three points mostly determine the sphere's position. (O)
- Use the dialog buttons to save or finish the annotation. (O)
- The spheroid annotation file is automatically saved next to the original image. (A)

##### *2. Marker Image Creator 3D (CMA)*

- Load the image. (O)
- Select nucleus (DAPI) channel and parameters for the LoG filter and maximum finder. (O)
- Apply 'Median (3D)' filter plugin on DAPI channel. (A)
- For each LoG filter step configured:
  - Apply LoG filter (by means of the LoG3D plugin [1]) on the filtered DAPI channel. (A)
  - Invert the LoG image to get proper maximums. (A)

  - Multiply by the XYRadius input parameter to normalize values for nucleus size. (A)
- If multiple LoG steps were taken, combine the results by taking the maximum value for every pixel position out of all the LoG filter images. (A)
- Apply an optimized variant of the 'Find Maximum…' plugin on the single or combined LoG filter image. (A)
- Create a new, black 3D stack with the sizes of nucleus channel and for all maximum points found points set the pixel value at its coordinates to a one-step incrementing value (starting at 1, so 1, 2, 3, 4, etc.). (A)
- Save the created marker image and a text file containing the coordinates of the maximum points, the label given, and the maximum value itself. (A)

#### *3. Marker Controlled Watershed 3D (CMA)*

- Load the image and the marker image. (O)
- Identify the nucleus and actin channels. (O)
- Select the filter and threshold methods for the nucleus and for the actin channel. (O)
- Select the parameter to correct for decreasing intensity with depth. (O)
- For the nucleus first and actin channel second:
  - Filter the channel according to the option chosen. (A)
  - Find the threshold for every slice of the channel. Do not apply the thresholds. (A)
  - For every slice, find the actual threshold by averaging over a number (set by user parameter) of the neighboring slice thresholds. If slice numbers outside of the image are required, take the closest existing slice threshold instead. (A)
  - Apply the averaged threshold for every slice. (A)
  - For the actin channel only, add the nucleus channel thresholded image. This is to ensure that the entire cell is segmented, including the nucleus. (A)
  - Create a 3D distance map for the thresholded image (uses the '3D Distance Map' plugin [2]). (A)
  - Use the distance map and the marker image as input for the marker controlled watershed (uses the plugin of the same name [3]). (A)
- Save both segment images. (A)

#### *4. Feature Extraction 3D (CMA)*

- Load the original image and the two segmented images (nucleus and actin). (O)
- Identify the nucleus and actin channels and also select any other channels to measure or alternative channel measurements. (O)
- Select any post-processing steps. (O)
- Load the marker coordinates file as produced by the Marker Image Creator 3D from the default location. (A)
- By means of the label identity in the marker coordinates file and the nucleus and actin segment images, create a list of nucleus and cell segment pairs. (A)

- Measure a configured list of features on the nucleus and cell segments. (This uses the standard measurements of Fiji as well as the methods provided by the plugins of the 3D ImageJ Suite [2] and MorphoLibJ [3]). (A)
- For every nucleus marker, if its coordinates are to close the any image edge (X, Y and Z) as configured by the user, flag the nucleus/cell segment pair as excluded by post-processing. (A)
- Load the spheroid annotation file and for each nucleus marker calculate the distance to the edge and the centre of the annotated spheroid. (A)
- For every additional channel or alternative measurement method measure the intensity of the following according to the measurement method:
    - Nuclear: Measure the intensity at all the coordinates of the nucleus segment. (A)
    - Nuclear centre: Measure the intensity within a radius of 3 around the nucleus marker. (A)
    - Cell: Measure the intensity at all the coordinates of the cell segment. (A)
    - Cell without the nucleus: Measure the intensity at all the coordinates of the cell segment excluding the coordinates of the matching nucleus segment. (A)
    - Nucleus surrounding:
        - Duplicate the cell segmentation image as a binary mask. (A)
        - Erode the masked segments. (A)
        - Measure the average intensity in a band of 2 pixels wide around the border of the nucleus segment. The coordinates measured should fall within the eroded mask, but not within any (other or same) nucleus segment. (A)
- For every cell segment that has not been excluded by post-processing, determine the migration mode:
    - Determine if any other cell segments are adjacent to this segment at any point; i.e. the minimum distance between any of the pixels of this segment and any pixel of another segment is 1. (A)
    - If no adjacency is found, the cell segment is considered a single cell. (A)
    - If any adjacency is found, determine the number of cell segments grouped via adjacency links to this cell segment. (A)
    - If the number is two, this cell is part of a paired migration group. (A)
    - If the number is greater than two it is considered a multi-cell migration group. (A)
    - The largest multi-cell migration group is determined to be the spheroid itself. (A)
- Create an image identifying the cell migration groups and an image identifying the migration mode per cell segment. Also create a duplicate image of the nucleus channel and draw the nucleus markers and the outlines of the nucleus segments on it (the latter in a separate color for excluded nuclei). (A)
- Create data files containing the measured features per nucleus and per cell and a file co
- containing a summary of the values over the entire image. (A)

*References:*

[1] D. Sage, F.R. Neumann, F. Hediger, S.M. Gasser, M. Unser, "Automatic Tracking of Individual Fluorescence Particles: Application to the Study of Chromosome Dynamics," IEEE Transactions on Image Processing, vol. 14, no. 9, pp. 1372-1383, September 2005

[2] J. Ollion, J. Cochennec, F. Loll, C. Escudé, T. Boudier. TANGO: A Generic Tool for High-throughput 3D Image Analysis for Studying Nuclear Organization. *Bioinformatics* 2013 Jul 15;29(14):1840-1
[3] Legland, D.; Arganda-Carreras, I. & Andrey, P. "MorphoLibJ: integrated library and plugins for mathematical morphology with ImageJ", Bioinformatics (Oxford Univ Press) 32(22): 3532-3534, 2016
